# Control of retrotransposon-driven activation of the interferon response by the double-stranded RNA binding protein DGCR8

**DOI:** 10.1101/2025.05.28.656609

**Authors:** A Gázquez-Gutiérrez, P Chin, G Peris, K Gordon, PG Marchante, J Witteveldt, P Tristán-Ramos, JO Rouvière, LI Knol, A Ivens, L López-Onieva, A Estevez, W Garland, TH Jensen, SR Heras, S Macias

**Author notes:** Correspondence should be addressed to SM or SRH.

## Abstract

The type I interferon (IFN) response is the main innate immune pathway against viruses in mammals. This pathway must be tightly regulated to prevent viral spread while avoiding excessive immune responses. Here, we show that inactivation of the double-stranded RNA (dsRNA)-binding protein DGCR8 unleashes the IFN response in human cells. We demonstrate that DGCR8 restricts the accumulation of endogenous dsRNA originating from protein-coding mRNAs that harbour transposable elements (TEs), primarily Alu. We propose that DGCR8 binding to TE-rich mRNAs is essential to resolve dsRNA structures, and in its absence, accumulated dsRNA signals through the RIG-I-like signalling pathway triggering the IFN response. This mechanism is relevant to conditions where DGCR8 expression levels are altered, including the 22q11.2 deletion syndrome (22qDS). Supporting this, we show that 22qDS-derived cells exhibit an exacerbated type I IFN response which inversely correlated with DGCR8 levels. All these together demonstrate the importance of suppressing endogenous TE-dsRNA accumulation to prevent unwanted immune activation and associated disease pathogenesis.

## Introduction

In mammals, the type I interferon (IFN) response constitutes the main innate immune response against viruses. Typically, the first step of the IFN response is the recognition of the viral nucleic acids. In the cytoplasm, this is driven by two main pathways: the cyclic GMP-AMP synthase (cGAS) and its downstream signalling effector stimulator of interferon genes (STING), which sense viral DNAs; and the retinoic acid-inducible gene I (RIG-I)-like receptor (RLR) signalling pathway, which recognises viral double-stranded RNA (dsRNA) and uncapped transcripts. These sensors initiate signalling cascades that converge in the activation of the TANK-binding kinase 1 (TBK1) and IkappaB kinases (IKKs) leading to the phosphorylation and nuclear translocation of the interferon regulatory factor 3 and 7 (IRF3/7), and NfkB^1,2^. These transcription factors induce the expression of type I IFNs and proinflammatory cytokines. Upon secretion, IFNs stimulate the Janus Kinases (JAKs) and the Signal Transducer and Activator of Transcription (STATs) signalling pathway, both in the infected and neighbouring cells. This results in the expression of hundreds of interferon-stimulated genes (ISGs) with both antiviral and inflammatory activities^3^.

In a subgroup of autoimmune diseases, the IFN response is constitutively activated in the absence of viral infection. These diseases, named type I interferonopathies, are generally caused by mutations in genes involved in RNA or DNA metabolism, which result in aberrant IFN activation by the recognition of self-nucleic acids^4^. Transposable elements (TE) have emerged as a significant source of endogenous immunostimulatory RNA and DNA, but it is still unclear if the nature of the pathogenic TEs responsible for triggering the IFN responses are conserved. Almost half of the human genome consists of retrotransposons, a specific class of TEs that propagate through a mechanism involving reverse transcription of the RNA intermediate to generate a new DNA copy of the element^5^. Based on their structure and mobilisation mechanism, retrotransposons are divided into two classes, those that contain long terminal repeats (LTR), and non-LTR retrotransposons. Although a significant fraction of TEs is transcribed, only a limited number of individual non-LTR retrotransposon copies retain the ability to mobilise in humans^6^. Active elements belong to the Long Interspersed Nuclear Element-1 (LINE-1) and the Short Interspersed Nuclear Element (SINE) families, including Alu and SVA elements^6^. The inactivation of genes involved in epigenetic silencing mechanisms of TEs, such as DNA methylation^7,8^ or the HUSH complex^9^, or interfering with the normal function of the double-stranded RNA binding proteins, such as ADAR1^10^ or MDA5^11^, result in TE-derived nucleic acid accumulation and IFN activation. Although many of these studies have attributed a role for TEs in the stimulation of the IFN response, it is usually not investigated if TE-derived RNAs actually originate from TE promoters, or whether they result from transcription of overlapping genes. This is an important aspect to study considering that most TE-derived RNAs originate from co-transcription hosted within mRNAs or by pervasive transcription^12,13^.

DGCR8 is a nuclear dsRNA-binding protein characterised for its essential role in microRNA (miRNA) biogenesis. Together with Drosha, DGCR8 forms the Microprocessor complex which cleaves primary miRNA precursors^14^. However, DGCR8 has been reported to play non-canonical functions independent of miRNA processing ^15,16^. Amongst these, DGCR8 has been shown to maintain heterochromatin organisation repressing TE expression^17^ and to bind TE-derived mRNAs restricting their mobilisation^18^.

Given the roles of dsRNA-binding proteins in controlling the type I IFN response, we aimed to address if DGCR8 could also regulate the IFN response in humans and if this ability was relevant to diseases where DGCR8 function is impaired. To this end, we generated *DGCR8* knockout human cell lines and found that in the absence of DGCR8, cells have an increased ability to mount an IFN response. We also observed that *DGCR8*-deficient cells exhibit a constitutive activation of the RIG-I-like receptor signalling pathway due to the accumulation of endogenous dsRNA. Such dsRNAs are enriched in retrotransposons embedded within protein-coding mRNAs, rather than self-expressed TE copies. We propose that this function of DGCR8 is independent of Microprocessor activity and is instead mediated by its widespread ability to bind dsRNA and resolve these structures through interaction with the RNA helicase DHX9. We also show that human diseases characterised by altered DGCR8 levels, including the 22q11.2 deletion syndrome (22qDS), where one copy of the DGCR8 gene is absent, also display an aberrant IFN response. Fibroblasts derived from 22qDS patients exhibit an exacerbated type I IFN response compared to cells from healthy individuals. This is conserved across all the studied patient cohorts and cell types, including blood samples and neuron-derived iPSCs from patients. These results support a non-canonical function for the dsRNA-binding protein DGCR8 in controlling the balance between endogenous dsRNA accumulation and immune sensing and activation.

## Material and methods

### Cell lines

All cell lines were grown at 37 °C and 5% CO_2_. Human PA-1 cells were cultured in Gibco MEM, 20% heat-inactivated foetal bovine serum, 100 U/mL penicillin-streptomycin and 0.1 mM Non-Essential Amino Acids (Gibco). HeLa cells were cultured in Dulbecco’s Modified Eagle Medium (DMEM) High Glucose (Sigma), 10% foetal bovine serum and 100 U/mL penicillin-streptomycin. Mouse microglial BV-2 cells were grown in Dulbecco’s Modified Eagle Medium (DMEM) High Glucose (Sigma), supplemented with 10% FBS, 2 mM L-glutamine (Gibco) and 100 U/mL penicillin-streptomycin.

E14TG2a mouse ESCs (male genotype, XY) and descended mTurboID-DGCR8 cell line were cultured on 0.2% gelatin-coated plates in 2i/LIF containing medium 1:1 mix of Neurobasal (Gibco) and DMEM/F12 (Gibco) supplemented with 100 U/ml penicillin-streptomycin (Gibco), 2 mM Glutamax (Gibco), 50 mM β-mercaptoethanol (Gibco), 0.1 mM Non-Essential Amino Acids (Gibco), 1 mM Sodium Pyruvate (Gibco), 0.5x N2 supplement (Gibco), 0.5x B27 supplement (Gibco), 3 mM GSK3i (CHIR99021, Sigma-Aldrich), 1 mM MEKi (PD0325901, STEMCELL Technologies) and Leukemia Inhibitory Factor (LIF, produced in house). CRISPR/Cas9-mediated genomic knock-ins of N-terminal 2xHA-mTurbo at the mouse DGCR8 locus was carried out using homology-dependent repair (HDR) donor vectors. Plasmids were generated containing specific 5’ and 3’ homology arms (∼ 500 bp) amplified from wildtype mouse ES cell genomic DNA and cloned into pGNT backbones. Tagging donor plasmids comprised of a 5’ DGCR8 homology arm-[HYG/PURO resistance]-P2A-[2xHA]-mTurbo-3’DGCR8 homology arm. sgRNA targeting the 5’ UTR of mouse DGCR8 was cloned into pSLCas(BB)-2A-GFP vectors (pX458 Addgene #48138) as previously described^19^. Cells were co-transfected using X-tremeGENE HP DNA Transfection Reagent (Roche) with 2 HDR donor plasmids harbouring distinct selection markers along with a sgRNA/Cas9 vector in a 1:1:1 ratio. Cells were selected for ∼ 7 days before picking and expanding single cell colonies. Clones were verified by western blot analysis and genotyping.

Primary fibroblast from 22qDS patients were purchased from Coriell Institute for Medical Research (GM07215F, GM02944) and control line from ATCC (CRL2429). Both were cultured in Gibco MEM, 15% heat-inactivated foetal bovine serum, 100 U/mL penicillin-streptomycin and 0.1 mM Non-Essential Amino Acids (Gibco). To achieve cellular immortalization, retroviral particles encoding the human telomerase reverse transcriptase (hTERT) were generated by transfecting Phoenix helper-free packaging cells with the pBABE-hygro-hTERT plasmid (Addgene plasmid #1773) using FuGENE HD transfection reagent (Promega). Human primary fibroblasts (GM07215F and CRL2429) at 70% confluency were transduced with viral supernatants supplemented with 5 µg/mL Polybrene (Sigma-Aldrich) for 24 hours. Seventy-two hours after transduction, cells were selected with 50 µg/mL hygromycin B (Thermo Fisher Scientific).

Inactivation of DGCR8 in PA-1 cells was performed by CRISPR/Cas9 against exon 2 as described in Colomer-Boronat et al. 2025^20^. For complete inactivation (knock-out), two clones were selected for further studies and named KO1 and KO2. One clone was selected for partial *DGCR8* expression inactivation as heterozygous (HET). Karyotyping for PA-1 was performed at the Andalusian Biobank (Granada). STR (Short tandem repeat) analysis was carried out at the Genomic Unit (Genyo, Granada).

### Cell proliferation

For the proliferation assay, xCELLigence RTCA MP (Agilent Technologies) was used. PA-1 WT, *DGCR8* KO1 and KO2 cells were seeded at 2×10^3^ cells per well in 96-wells plate. Measurements were made every hour for a week and media was replaced after four days. GraphPad Prism 8.3 software was used for analysis and graph preparation.

### DNA and RNA viral mimics, interferon inhibition and stimulation, retrotranscriptase inhibitors and chemotherapeutic agents

Cells were transfected at 70% confluency, in six-well plates, and transfected with 5 μl of Invitrogen Lipofectamine 2000 Transfection Reagent and 1 μg of Y-shaped-DNA cGAS agonist (G3-YSD) (#tlrl-ydna, Invivogen) or dsRNA analogue polyinosinic:polycytidylic acid (poly(I:C)) (HMW #tlrl-pic, Invivogen). Cells were transfected for 6 and 4 hours respectively before media replacement. After G3-YSD or poly(I:C) treatment, RNA was extracted using Trizol (Thermo Fisher) at 16 hours post-transfection. For interferon inhibition, Ruxolinitib (#tlrl-rux, Invivogen) and BX-795 (#tlrl-bx7, Invivogen) were added to the media, at 50 μM and 8 μM respectively, for 6 hours before media replacement. RNA was extracted at 16 hours post-addition. For stimulation with exogenous IFN, cells were treated with 200 U/ml of exogenous human IFN-*β* (HZ-1298m Proteintech) for 6 h and harvested in Trizol for RNA extraction. Retrotranscriptase inhibitors (RTis) used to treat HIV/AIDS and inhibit L1 reverse transcriptase^21^, emtricitabine and lamivudine, were obtained from Hospital Clínico San Cecilio, Granada, Spain. Cells were treated with 25 μM of lamivudine or emtricitabine for six days, replacing media with fresh RTi every two days. Cells were treated with 7 μM of etoposide (2200S, Cell Signalling) for 0h, 8h and 24h. Each experiment was performed in parallel plates containing three biological replicates for each condition, where cells were fixed for immunofluorescence against γH2AX, and additional replicates were used for RNA extraction.

### Viral infection

Stocks of RSV strain A2 (kindly provided by Dr. Sarah Ressel, University of Edinburgh) and Influenza A virus strain PR8 (kindly provided by Prof. Paul Digard, University of Edinburgh) were grown in HEp-2 and MDCK cells. RSV susceptibility was determined using the 50% tissue culture infectious dose (TCID_50_) assay in 96-well format. For this, cells at approximately 50% confluency were infected in serum-free medium with seven serial dilutions, using six wells per dilution. After 2 hours, medium was replaced with serum-containing medium. Cytopathic effects were recorded 48 hours post-infection and TCID_50_ values were calculated using the Spearman and Kärber algorithm. For interferon inhibition, cells were incubated with 10 μM of ruxolitinib (ab141356, Abcam) for 1 hour pre-infection. Infections were performed as before in serum-free medium containing 10 μM of ruxolitinib. After 2 h, the virus inoculum was removed and complete medium supplemented with 10 μM of ruxolitinib was added. Influenza A virus replication was determined by infecting cells at the same multiplicity of infection (MOI) in serum-free medium supplemented with 2 µg/ml TPCK-treated trypsin for 45 minutes. After replacement of the inoculum with fresh serum-containing medium, the cells were incubated for 48 hours, lysed in Trizol before RNA isolation.

### Immunofluorescence

PA-1 WT cells and PA-1 *DGCR8* KO cells were seeded at 2×10^5^ cells/well and 4×10^5^ cells/well on a glass coverslip on a 6-well plate. For actinomycin treatment, actinomycin D (A1410 Sigma-Aldrich) was diluted in DMSO and supplemented to the cell media at 0.5 μg/ml for 2 hours. Cells were fixed with 2% paraformaldehyde in PBS for 20 minutes and permeabilised with 0.4% Triton X-100 in MiliQ water for 5 minutes. Wells were washed twice in PBS and twice in wash buffer (PBS, 0.5% Tween 20, 0.1% BSA). For cells subjected to RNase III treatment (ShortCut RNase III #M0245 NEB), coverslips were incubated with 40 U/ml RNase III in PBS supplemented with 5 mM MgCl_2_ for 30 minutes at 37°C and then washed. Cells were blocked with blocking buffer (PBS, 0.5% Tween 20, 2.5% BSA, 10% goat serum) for 30 minutes. Coverslips were incubated with J2 primary antibody (10010200, Nordic) or isotype (3878, Santa Cruz) at 5.625 μg/ml in blocking buffer for two hours at room temperature before washing three times in washing buffer. Coverslips were incubated in secondary antibody (AlexaFluor488 donkey anti-mouse, A-21202, Thermo Fisher) at 4 μg/ml in blocking buffer for 45 minutes in the dark at room temperature. Coverslips were washed twice in PBS and washing buffer and nuclei were stained using ProLong Gold antifade (P10144, Invitrogen). Zeiss LSM 710 Confocal Microscopy with a Plan-Apochromat 63x/1.40 Oil DIC M27 was used for imaging. Image J^22^ was used for quantification of 150 cells.

For γH2AX staining, coverslips were incubated with 10 μg/ml of primary antibody (NB100-74435, Novus Bio) followed by secondary antibody Alexa Fluor 555 goat anti-mouse IgG (A21424, Invitrogen) at 4 μg/ml.

### Flow cytometry

PA-1 WT or KO cells were detached and filter through a 70 μM cell strainer and 2×10^6^ cells per condition were split into flow cytometry tubes. Cells were blocked for 10 minutes in Fc blocking (101319, Biolegend) 5 µg/ml in FACS buffer (PBS, 0.5% BSA, 2 mM EDTA) and fixed and permeabilised using BD Cytofix/Cytoperm solution (554714, BD). Cells subjected to RNase III treatment (M0245, NEB) were incubated with 40 U/ml RNase III in PBS supplemented with 5 mM MgCl_2_ for 30 minutes at 37°C. Cells were washed using BD Perm/Wash buffer (554714, BD) and incubated with primary antibody (10010200, Nordic) or isotype (3878, Santa Cruz) at 1.875 μg/ml for one hour in BD Perm/Wash buffer. Cells were washed and incubated in secondary antibody (Alexa Fluor 488 donkey anti-mouse, #A-21202, ThermoFisher) at 2 μg/ml in BD Perm/Wash buffer, for one hour. Cells acquisition was performed using a FACSVerse cytometer. Due to considerable differences in cell size, gating for J2 positive cells was adjusted for each cell line according to the isotype control signal. FlowJo Software was used for quantification and preparation of figures.

### Western Blotting

Cells were lysed in RIPA buffer supplemented with 1x EDTA-free Protease Inhibitor cocktail (11873580001, Sigma-Aldrich), phosphatase inhibitor cocktail 2 (P7526, Sigma-Aldrich) and phosphatase inhibitor cocktail 3 (P0044, Sigma-Aldrich), 1mM PMSF and 35nM β-mercaptoethanol. Pierce BCA Protein Assay Kit (Thermo Fisher Scientific) was used for protein quantification. Lysates were subjected to SDS-PAGE electrophoresis and transferred to PVDF membranes using Trans-Blot Turbo Transfer System (Bio-Rad). Membranes were blocked in 5% non-fat dried milk or BSA in TBS-T (0.1% Tween-20 in TBS). Primary antibodies were incubated overnight at 4°C and secondary antibodies for one hour at room temperature. Images were acquired using an ImageQuant LAS4000 or Chemidoc Imaging System (Bio-Rad), fluorescent signal was quantified using ImageJ and Chemidoc Imaging softwares. Antibodies used include anti-DGCR8 (1:1000, ab90579, Abcam), anti-Drosha (1:1000, NBP1-03349, NovusBio), anti-pEIF2α (1:1000, 3398, Cell Signalling), anti-IFIT3 (1:500, ab76818, Abcam), anti-RIG-I (1:1000, 3743, Cell Signalling), anti-DHX9 (1:1000, 70998 Cell Signalling), anti-ZCCHC8 (1:1000, A301-806 Thermo Fisher Scientific), anti-Ago2 (1:1000, 2897S Cell Signalling), anti-α-tubulin (1:1000, sc-23948, Santa Cruz Biotechnology), anti-vinculin (1:100000, V9131 Sigma Aldrich) anti-β-actin (1:15000, A1978, Sigma-Aldrich), anti-GAPDH (1:1000, CB1001, Sigma Aldrich). As secondary antibodies, anti-rabbit-HRP (1:1000, 7074S, Cell Signaling), anti-mouse-HRP (1:1000, 7076S, Cell Signaling) and anti-goat-HRP (1:10000, 305-035-003, Jackson ImmunoResearch).

### RNA extraction and RT-qPCR

RNA was extracted using TRizol (15596026, Invitrogen) or TRizol LS (10296010, Invitrogen) following manufacture’s instruction. RNA was subjected to initial DNase treatment using RQ1 DNase (M6101, Promega) for one hour at 37 °C, followed by phenol/chloroform purification and RNA quantification. One µg of RNA was further treated with DNase I (18068015, Invitrogen), and cDNA was synthesised using High-Capacity cDNA Reverse Transcription Kit (4368814, Applied Biosystems). cDNA was diluted 1:4 and used for qPCR (GoTaq qPCR Mix, A6002 Promega). Primers amplifying human *GAPDH, ACTB, RN7SK,* and *18S* rRNA were used as normalisers and gene expression levels were quantified using the second derivative method. Primers used are listed in **Supplementary Table S1**.

### Protein immunoprecipitation (IP) and proximity labelling (PL)

For whole cell extract IPs, 2×10^7^ cells/IP were extracted in HT150 extraction buffer (20 mM HEPES pH 7.4, 150 mM NaCl, 0.5% v/v Triton X-100) freshly supplemented with protease inhibitors. Lysates were sheared by sonication (3 x 5 s, amplitude 2) and cleared by centrifugation at 18,000 rcf for 20 min. Clarified lysates were incubated with FLAG antibody (F3165, Sigma) overnight at 4 °C with Protein G Dynabeads (10003D, ThermoFisher). Beads were washed 3 times with HT150 extraction buffer, transferring beads to a fresh tube on the final wash. Proteins were eluted by boiling in 1X NuPAGE loading buffer (Invitrogen) for 5 min. Supernatants were mixed with 10X Reducing Agent (Invitrogen) and denatured for a further 5 min at 95 °C before proceeding with western blotting analysis. For Benzonase treated IPs, samples were extracted and washed in HTM200 buffer (20 mM HEPES pH 7.4, 200 mM NaCl, 1 mM MgCl2, 0.5% Triton X-100). After the final washing step, beads were resuspended in HTM200 buffer and either mock or treated with 250 units of Benzonase (Sigma) at 25 °C for 20 min in a thermomixer at 1000 rpm. Beads were pelleted and supernatants were collected for Benzonase elution samples. The beads were washed 2x in HTM200 before elution of bound proteins using the standard protocol described above. For proximity labelling, cells were treated with 50 µM of biotin for 24h and HT150 extracted lysates were incubated with Dynabeads MyOne Streptavidin C1 (65001, Invitrogen) overnight to enrich for biotinylated proteins. Beads were treated as above and samples were analysed by western blotting.

### Proteomic mass spectrometry

PA-1 WT and *DGCR8* KO2 cells (n=3) were pelleted, washed three times with DPBS, and stored at –80 °C until further processing and analysis, following the protocol described in Ryan et al. 2021^23^. Briefly, 100 µg pellets were lysed, reduced, and alkylated in 100 µL of 6 M guanidine hydrochloride, 200 mM Tris-HCl pH 8.5, 1 mM TCEP, and 1.5 mM chloroacetamide. Samples were sonicated, heated at 95 °C for 5 min to complete reduction, and centrifuged. Proteins were digested overnight with sequencing-grade trypsin. Following digestion, samples were acidified, and approximately 20 µg of peptides were desalted using StageTips. Eluted peptides were freeze-dried, resuspended in 0.1% TFA, and quantified using a NanoDrop spectrophotometer. Samples were then diluted to 200 ng/µL for liquid chromatography–tandem mass spectrometry (LC-MS/MS) analysis. Peptide mixtures (5 µL per sample, corresponding to 1 µg) were injected into an Aurora column (IonOpticks) on an UltiMate 3000 HPLC system (Thermo Fisher Scientific) coupled to an Orbitrap Fusion Lumos Tribrid mass spectrometer (Thermo Fisher Scientific). Peptides were separated with a 150-minute reverse-phase gradient from 3% to 40% acetonitrile at a flow rate of 400 nL/min and a voltage of 1500 V. Data were acquired with an MS resolution of 240,000, a 1-second cycle time, and MS/MS HCD fragmentation with analysis in the ion trap. Raw MS spectra were analysed with MaxQuant using label-free quantification (LFQ) against the UniProt Homo sapiens database, including all protein isoforms. A false discovery rate (FDR) of 1% was applied at both peptide and protein levels. All three replicates were grouped for statistical analysis. Contaminants and exact duplicate entries were removed prior to downstream analysis. Data were normalized, and log2 fold changes were calculated using the Limma R package^24^. Proteins were considered significantly differentially expressed with adjusted p-values < 0.05 and absolute log2FC > 0.5.

### RNA immunoprecipitation (RIP)

2 µg of DGCR8 antibody (ab90579, Abcam) per RIP was incubated overnight at 4 °C with rotation with pre-washed Dynabeads Protein G (10003D, ThermoFisher). 1×10^7^ cells were washed with ice-cold 1× PBS, scraped and transferred to a 1.5 ml tube. After centrifugation at 200 x *g* for 4 min, cells were resuspended in 200 µl of cold resuspension buffer (20 mM Tris pH 7.5), 150 mM NaCl, 1 mM EDTA, 1 mM EGTA) containing 1 U/µL RNAsin Plus (N2611, Promega) and lysed adding 800 µl of cold lysis buffer (1% Triton X-100, 20 mM Tris pH 7.5, 150 mM NaCl, 1 mM EDTA, 1 mM EGTA, 1 mM phenylmethyl-sulfonyl fluoride (PMSF, 329-98-6, Sigma), 1X complete EDTA-free Protease Inhibitor cocktail (04693159001, Merck)) and incubated for 20 min on ice. After centrifugation (10,000 x *g* for 10 min at 4 °C), 10 µL of RQ1 DNase (M6101, Promega) was added to the supernatant. Immunoprecipitation with antibody-coupled beads was performed at 4 °C with rotation for 3 hours. After five washes with lysis buffer, 10% of sample-beads were used for protein extraction and western blot by adding LDS sample buffer and DTT and heating the samples at 70 °C for 20 min. The 90% of sample beads were incubated with RQ1 DNase for 30 min for later RNA extraction with Trizol LS (11578616, ThermoFisher).

### J2 immunoprecipitation

Cell pellets from WT, *DGCR8* KO1 and KO2 were lysed in lysis buffer (10mM Tris pH 7.4, 150 mM NaCl, 0.075% IGEPAL, supplemented with 40U of RNase inhibitors (N2611, Promega), RQ1 DNase (M6101, Promega), 0.5 mM DTT, 0.2 mM PMSF, and cOmplete EDTA-free protease inhibitors (4693159001, Merck) on ice for 5 min, followed by centrifugation at 3,500 x g for 5 min. Supernatants were centrifuged again for 5 min at 13,000 rpms to remove any debris, and saved as cytoplasmic fraction. 50 µl of slurry Protein G Dynabeads (10003D, ThermoFisher) were washed 3 times with IP buffer (50 mM Tris pH 7.5, 150 mM NaCl, 1 mM EDTA, 1% Triton X-100), and coupled to 5 µg of anti-dsRNA monoclonal antibody (J2, RNT-SCI-10010200 Scicons) for 1 h at RT, followed by 3 washes with IP buffer to remove unbound antibody. Cytoplasmic supernatants were diluted in IP buffer, supplemented with 0.5 mM, 0.2 mM PMSF and Roche cOmplete EDTA-free protease inhibitors, and incubated with antibody-coupled beads for 4 hours at 4°C. Beads were washed 3 times with IP buffer, changing tubes after each wash. RNA was extracted from beads using Trizol LS and precipitated overnight with isopropanol. RNA quality was assessed by Bioanalyzer (Agilent) and sequenced by the Clinical Research Facility at the University of Edinburgh. Briefly, 5 ng of each IP-RNA sample (3 biological replicates for each cell line; WT, KO1 and KO2) was used for library preparation using the NEBNEXT Ultra II Directional RNA Library Prep Kit (E7760, NEB) and the NEBNext rRNA Depletion kit (Human/Mouse/Rat) (E6310, NEB). Sequencing was performed on the Illumina NextSeq 2000 platform using NextSeq 1000/2000 P2 Reagents (200 Cycles) v3 (20046812, Illumina Inc). 50 million pair-end reads were produced for each sample.

### Total RNA high-throughput sequencing

Total RNA from four biological replicates of PA-1 WT and *DGCR8* KO2 cells was extracted using miRNeasy Mini Kit (Qiagen) followed by on-column DNase digestion. Purified RNA was rRNA-depleted prior to sequencing (DNB-seq) by Beijing Genomics Institute (BGI).

### RNA-sequencing preprocessing and expression analysis

The following preprocessing pipeline was used for all sequencing methods (total RNA-seq and dsRNA-seq). First, FastQC v0.11.5^25^ was used to assess quality control metrics for paired end sequencing reads and BBDuk from the BBMap toolkit (https://sourceforge.net/projects/bbmap/) for sequencing adapter removal and read trimming. Next, RNA paired-end reads were aligned to GRCh38.p14 human genome assembly with STAR v2.7.6a^26^ and quantified with featureCounts v2.0.1^27^ using NCBI Annotation Release 102 and with option *--extraAttributes “gene_biotype“* for annotating sequence types. Differential expression analysis and count normalisation were performed with the R package DESeq2 v1.36^28^. Shrinkage of effect size was performed on DESeq2 results using the *apeglm* method through the function lfcShrink^29^. Volcano plots were generated with ggplot2 and heatmaps with R package ComplexHeatmap v2.4.3^30^.

### Read distribution

To analyse the distribution of reads for different features, such as protein coding, mobile elements or non-coding RNA, we followed the guidelines recommended in Teissandier et al. 2019^31^ for alignment with STAR: *--runThreadN 4 --outSAMtype BAM SortedByCoordinate --runMode alignReads --outFilterMultimapNma× 1000 -- outSAMmultNmax 1 --outFilterMismatchNmax 3 --outMultimapperOrder Random -- winAnchorMultimapNma× 1000 --alignEndsType EndToEnd --alignIntronMax 1 -- alignMatesGapMax 350.* To quantify repetitive elements with featureCounts, we reported randomly one position per read: *-M -s 0 -p*. Reads aligned to features were normalised by the number of aligned reads. Stacked bar plots were generated using ggplot2.

### Small RNA-sequencing analyses

Four biological replicates of WT and *DGCR8* KO2 PA-1 cells were sequenced by small RNA sequencing (Beijing Genomics Institute, BGI). Small RNA-seq libraries were generated using unique molecular identifiers (UMIs) and sequenced using DNBseq^25^. All sequencing reads were pre-filtered by BGI, with adapters, low quality reads and contaminants being removed, and data was returned in compressed FASTQ format. Quality control for all three datasets by FastQC v0.11.5^25^ showed that reads were of high quality and no further quality filtering was required.

For each library, identical small RNA-seq reads were collapsed and only reads that were present more than once were included in further analysis. Mature miRNAs were identified and counted using miRDeep2, using the quantification function with human hairpin and mature miRNA sequence files from miRbase v22.1 and allowing no mismatches^32–34^. In addition, the small RNA samples were analysed with blastn v2.6.0+ using the commands *-num_threads 20 -max_target_seqs 1 -max_hsps 1 1 -outfmt ‘6 std qseq sseq’ -task blastn -word_size 6 -dust no* in order to identify small RNAs^35^. DEseq2 was used for statistical analysis using *apeglm* as log fold-change shrinkage model, using the miRDeep2 counts as input^28,29^. Instead of using DESeq2’s built-in normalisation, the counts were normalised to the amount of tRNAs, relative to WT sample 1, identified in each sample in the blastn analysis using the *normalizationFactors* option of DESeq2.

Iteration of statistical analysis occurred eliminating the miRNAs with the overall lowest expression until all miRNAs analysed passed the independent filtering threshold with alpha = 0.1. In doing this, overall lowly expressed miRNAs were filtered up to the point that no miRNAs were removed from statistical analysis due to low read count, resulting in expression data for 700 mature miRNAs. MiRNAs with an FDR lower than 0.05 were considered significantly differentially expressed.

### Transposable element expression analyses

The Software for Quantifying Interspersed Repeat Expression (SQuIRE) v0.9.9.92^36^ was used with default parameters to quantify transposable elements expression in all datasets, both at family and individual loci level. Expression analyses were carried out using DESeq2.

### Origin of TE expression

To determine whether the expression of TEs arises from the TE promoter or is driven by a nearby gene, we followed the pipeline used by Chang et al 2022^37^ with modifications. Using this approach, a TE is considered gene-dependent if it overlaps with an exon, UTR, or intron of an expressed protein coding gene (with at least 10 normalised counts at any sample in at least one clone). TEs are also considered gene-dependent if lying inside a lengthened 3’ UTR region: 3’ UTR were considered extended when transcripts assembly using StringTie v1.3.3b^38^ revealed regions beyond 3’ UTR annotation. Alternatively, a TE is considered self-expressed if it is located in an intergenic region or inside introns of non-expressed genes. To annotate TEs position relative to genomic features, ChIPseeker v1.39.0^39^ was used employing the priority: “5UTR” > “3UTR” > “Exon” > “Promoter” > “Intron” > “Downstream” > “Intergenic”. Then, scripts in Chang et al. 2022 pipeline^37^ were adapted to fit SQuIRE and a novel gene-dependent category was introduced, classifying as “antisense” those TEs expressed in intergenic or non-expressed introns in the opposite strand such TE is annotated.

### TE density over features

An in-house R script *TE_feature_density* was developed to compute the number and density of TEs in protein-coding gene features (gene, exons, introns, 5’ UTR, 3’ UTR, downstream). This script can be found in https://github.com/GuillePeris/TE_feature_density) and downloads both TE and gene annotations from Ensembl (using R packages biomartr v1.0.7^40^ and UCSCRepeatMasker v3.15.2 (https://www.repeatmasker.org)) or reads them from user files, and then intersects gene features with TEs coordinates using R package bedtoolsr v2.30.0-5^41^, a wrapper for bedtools^42^.

### High-throughput sequencing of RNA isolated by crosslinking immunoprecipitation (HITS-CLIP)

We re-analysed previously generated HITS-CLIP data from endogenous and T7-tagged, overexpressed DGCR8 immunoprecipitated in human HEK293T cells (GSE39086)^43^. Hg18 peak coordinates were converted to hg38 with liftOver^44^ and annotated with ChIPseeker v1.39.0^39^ relative to protein-coding features. Only peaks appearing in at least two out of the four samples were considered. To further filter background noise, gene expression from HEK293T was retrieved from Tchurikov et al. 2019^45^ (GEO repository under accession no. GSE130262) and only genes with TPMs over 1 were considered.

### 22qDS datasets analysis

Publicly available transcriptomic datasets for patients with 22q11.2 deletion syndrome were obtained from the Gene Expression Omnibus (GEO) and analysed. Raw RNA-seq data were downloaded and processed for two datasets: neurons derived from induced pluripotent stem cells (iPSCs) of patients (GSE46562, n = 8)^46^ and T cells isolated from peripheral blood (GSE184790, n = 13)^47^. For additional datasets, differential expression analysis results were extracted from published studies, including iPSCs at different stages of neuronal differentiation (human pluripotent stem cells, hPSCs; neural progenitor cells, NPCs; and neurons) (n = 20)^48^, cerebral cortical spheroids derived from patient iPSCs at different stages of differentiation (n = 15)^49^ and peripheral blood samples (n = 79)^50^.

For gene ontology analysis, genes from 22q11.2 deletion syndrome patient datasets were ranked based on their log2 fold change values, weighted by the negative logarithm of their p-values, and sorted in descending order. GSEA was performed using the gseGO function from the R package clusterProfiler v4.12.1^51^. Multiple testing correction was applied using the Benjamini-Hochberg method with a significance threshold of 0.05. To reduce redundancy, enriched GO-BP terms with a similarity score above 0.7 were merged, retaining the most significant terms. Enrichment scores were used to classify terms as “activated” or “suppressed”. Dot plots were generated to visualize the top GO-BP terms by lowest adjusted p-values.

For heatmaps and correlations, a list of annotated interferon-stimulated genes (n=378) was used^52^.

### 3’ end RNA sequencing analysis

Published 3’end-seq from WT Hela RNA samples treated with *E. coli* pA polymerase^53^ (GSE179106) were used. 3’end RNA-seq data is represented as aggregate plots generated using custom R code showing the mean log2-transformed signal coverage, with a 90% confidence interval. For SINEs, signals were plotted 300 bp upstream and downstream of the SINE of interest, and the transcript body area (TSS to TES) was normalised to 300bp. For miRNAs, signals were plotted 25 bp upstream and downstream of the miRNA of interest. The number of aggregated transcripts (n) is indicated below the plots. The 10^−1^ multiplicated factor displayed at the top left of the plot should be applied to the whole plot scale.

## Results

### DGCR8 protects from constitutive IFN signalling

To understand the role of DGCR8 in controlling the IFN response, we inactivated the *DGCR8* gene using the CRISPR/Cas9 nickase editing system in human embryonic teratocarcinoma PA-1 cells, which are a stable diploid cell line^54^. Two different *DGCR8* knockout clones (KO1 and KO2) were selected for further studies. Inactivation of both *DGCR8* alleles abolished DGCR8 protein expression (**Fig 1A**), and resulted in reduced DROSHA protein levels, as its stability requires interaction with DGCR8 (**Fig 1B**)^55^. At the cellular level, *DGCR8* KO cells exhibited reduced proliferation rate (**Supp Fig 1A**) and maintained a stable diploid karyotype (**Supp Fig 1B-D**).

**FIGURE 1.**
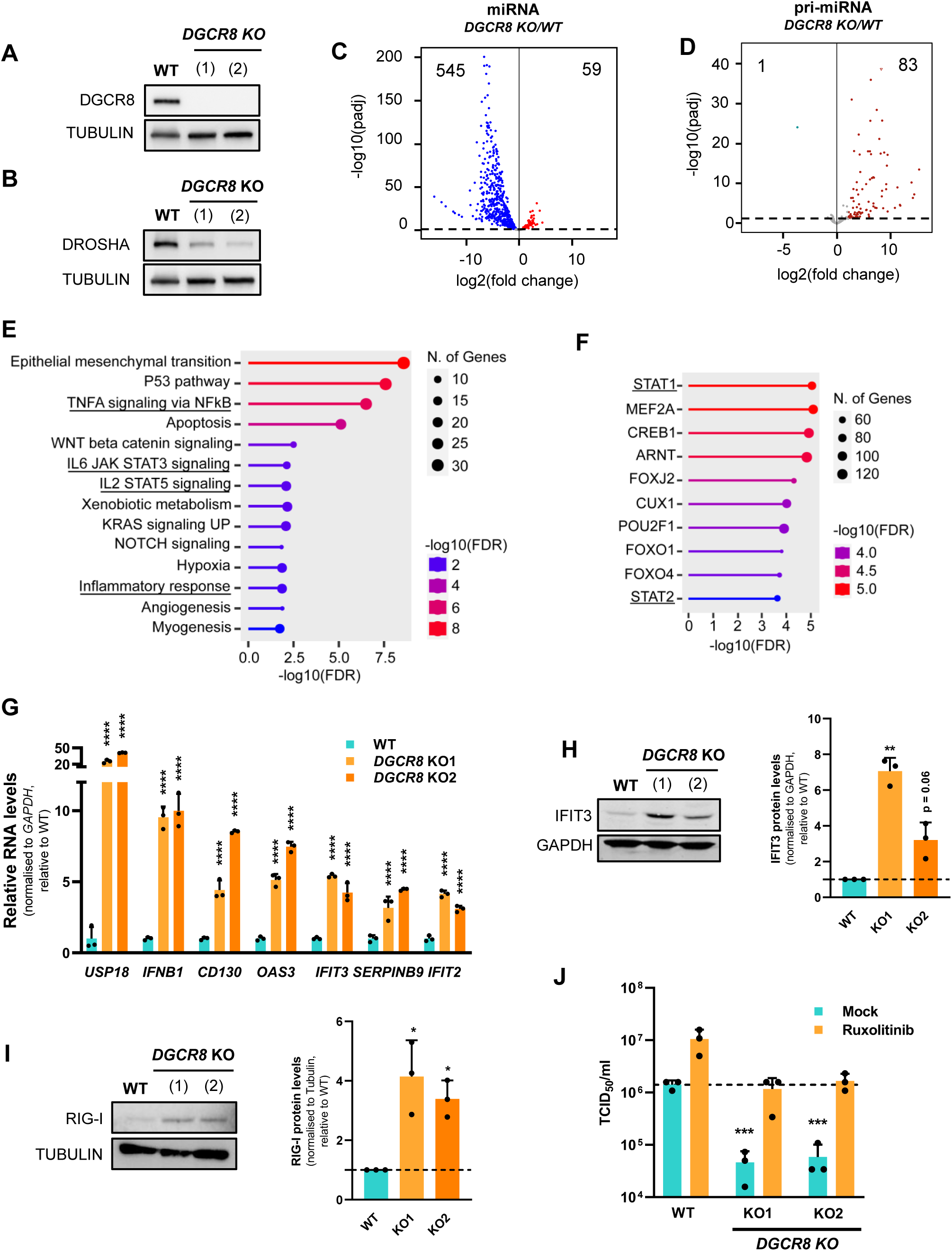
*DGCR8* KO cells show a basal and functional IFN activation. a. Western blot analyses of DGCR8 protein levels in PA-1 WT cells vs two *DGCR8* KO clones. Tubulin serves as loading control. b. Western blot analyses of Drosha protein levels in PA-1 WT cells vs two *DGCR8* KO clones. Tubulin serves as loading control. c. Volcano plot for differential miRNA expression of *DGCR8* KO2 vs WT cells using small-RNA sequencing. Blue represents significantly downregulated miRNAs (log2FC < −1 and p-adj < 0.05), red represents significantly upregulated (log2FC > 1 and p-adj < 0.05). d. Volcano plot for differential pri-miRNA expression in *DGCR8* KO2 vs WT, using total RNA-sequencing. Blue represents significantly downregulated pri-miRNAs (log2FC < −1 and p-adj < 0.05), red represents significantly upregulated (log2FC > 1 and p-adj < 0.05). e. Hallmark gene sets obtained with upregulated genes in *DGCR8* KO2 cells vs WT (log2FC > 0.6 and p-adj < 0.05). f. Transcription Factor (TF) targets network (ENCODE) obtained using the upregulated genes in total RNA-Seq (log2FC > 0.6 and p-adj < 0.05) in *DGCR8* KO2 vs WT. g. RT-qPCR analysis of ISGs and *IFNB1* in WT and two *DGCR8* KO clones in unstimulated conditions. *GAPDH* serves as a normaliser, and quantification is expressed relative to WT levels. h. Western blot analyses of the ISG IFIT3 in PA-1 WT and two *DGCR8* KO clones. GAPDH is used as loading control. On the right, quantification of three independent biological replicates. The dashed line represents the WT mean. i. Western blot analyses of the ISG RIG-I in PA-1 WT and two *DGCR8* KO clones. Tubulin serves as loading control. On the right, quantification of three independent biological replicates. The dashed line represents the WT mean. j. Susceptibility (TCID_50_/ml) of WT and two *DGCR8* KO cell lines to respiratory syncytial virus (RSV) infection in the presence and absence of ruxolitinib. The dashed line represents the WT mean. Bars represent the average of at least three replicates. Error bars plot mean ± SD. *p < 0.05, **p < 0.01, ***p < 0.001, ****p < 0.0001. P-values are determined by one-way ANOVA followed by Dunnett multiple comparison post-hoc test.

At the molecular level, small RNA high-throughput sequencing analysis revealed a strong downregulation of most mature miRNAs in DGCR8 KO cells, as expected (**Fig 1C**), in addition to accumulation of primary miRNA precursors (pri-miRNAs), as shown by total RNA sequencing (**Fig 1D**). In addition to miRNAs, DGCR8 deficiency resulted in widespread dysregulation of gene expression as revealed by RNA-seq analysis (495 upregulated and 1216 downregulated genes, |log2FC| > 1 and p-adj < 0.05, **Supp Fig 2A**, **Supp Table S2**). Upregulated genes belonged to biological pathways related to inflammatory responses, including ‘TNFα signalling via NFkB’, unlike downregulated genes which were not enriched in these processes (**Fig 1E, Supp Fig 2B**). STAT1 and STAT2, both involved in IFN signalling, were amongst the most enriched transcription factors predicted to account for the observed differential gene expression (**Fig 1F**). To further explore IFN activation in these cells, we analysed the interferon-stimulated gene signature (ISGs) at the RNA level by using RT-qPCR and total RNA-seq (**Fig 1G** and **Supp Fig 2C**) and at the protein level, using western blot and whole-cell proteomics (**Fig 1H-I**, **Supp Fig 2D** and **Supp Table S3**). Together this confirmed that *DGCR8* KO cells displayed an upregulation of ISG expression.

**FIGURE 2.**
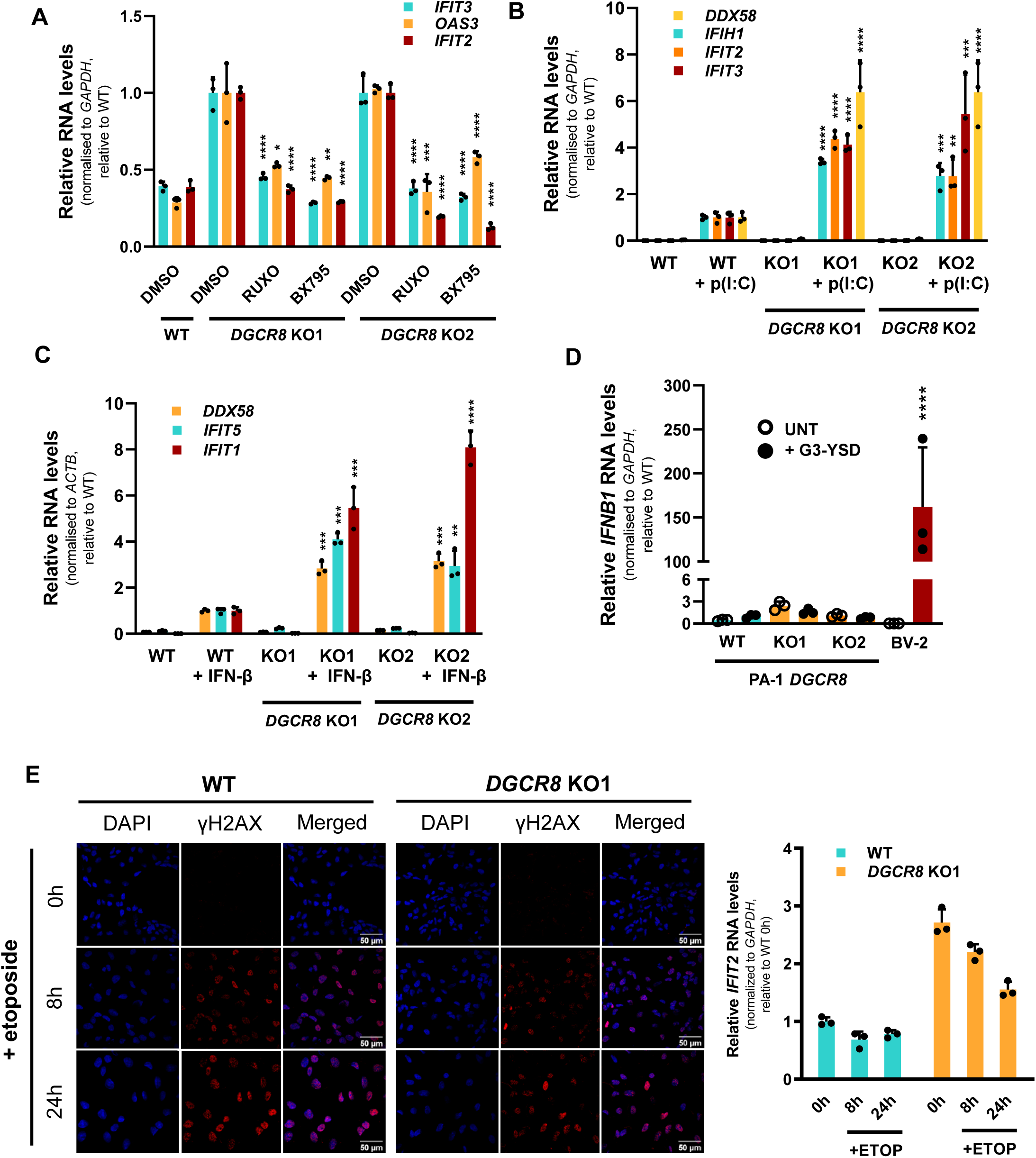
Basal IFN activation of *DGCR8* KO cells relies on the dsRNA-sensing pathway. a. RT-qPCR analysis of the ISGs *IFIT3*, *OAS3* and *IFIT2* in WT and two *DGCR8* KO clones treated with BX795, ruxolitinib (RUXO) or vehicle control (DMSO). *GAPDH* serves as a normaliser. ISG expression is represented relative to the average expression of each gene in the DMSO condition. b. RT-qPCR analysis of the ISGs *DDX58*, *IFIH1*, *IFIT2* and *IFIT3* in WT and two independent *DGCR8* KO clones transfected with poly(I:C). *GAPDH* serves as a normaliser. ISG expression is represented relative to the average expression of the WT+p(I:C) condition. c. RT-qPCR analysis of the ISGs *DDX58*, *IFIT5* and *IFIT1* in WT and two independent *DGCR8* KO clones after stimulation with exogenous IFN-β. *ACTB* serves as a normaliser. ISG expression is represented relative to the average expression of the WT+IFN-β condition. d. Expression levels of the *IFNB1* mRNA in WT and two *DGCR8* KO clones in after G3-YDNA transfection by RT-qPCR. BV-2 cells were used as a positive control for successful cGAS stimulation. *GAPDH* serves as normaliser. Expression is represented relative to the average expression of the WT+ G3-YDNA condition. e. WT and *DGCR8* KO1 cells were treated with etoposide for 0h, 8h or 24h. On the left, immunofluorescence of γH2AX which serves as a control of etoposide-induced DNA damage. DAPI is used for nuclear counterstain. Scale bar represents 50 µm. On the right, RT-qPCR analyses of *IFIT2* levels after etoposide treatment. *GAPDH* serves as normaliser. *IFIT2* levels are relative to 0h WT cells. Bars represent the average of three biological replicates. Error bars plot mean ± SD. *p < 0.05, **p < 0.01, ***p < 0.001, ****p < 0.0001. P-values are determined by one-way ANOVA followed by Dunnett multiple comparison post-hoc test.

We next wanted to assess whether the upregulation of ISGs expression was sufficient to lead to differential viral susceptibility of WT and *DGCR8* KO cells. To this end, cells were infected with respiratory syncytial virus (RSV), a negative-sense, single-stranded RNA virus. *DGCR8* KO cells were at least 10 times more resistant to the virus than WT cells (**Fig 1J**). To test whether the differences in viral infectivity were due to differential ISG expression, we treated cells with the JAK-STAT inhibitor, ruxolitinib, which blocks IFN signalling. This treatment increased the sensitivity of cells to RSV infection, with *DGCR8* KO cells reaching similar levels of susceptibility as WT cells. This indicates that differences in infectivity between WT and *DGCR8* KO cells are for the majority driven by IFN signalling (**Fig 1J**). These results demonstrate that the absence of DGCR8 leads to a sterile and constitutive activation of the type-I IFN response.

### ISG expression depends on the RIG-I-like receptor (RLR) signalling

The upregulated expression of ISGs in *DGCR8* KO cells could be the consequence of a global reduction in miRNA levels, as these are well-described post-transcriptional regulators of innate immune genes, including ISGs themselves or the IFN receptor^56–58^. Alternatively, ISG levels may be increased due to a constitutive activation of the IFN response. To test these possibilities directly, we blocked the IFN response at several steps of the pathway while measuring the expression of ISGs. Our initial attempt consisted of knocking down crucial signalling proteins of the IFN response using siRNAs and while WT cells could be efficiently depleted, *DGCR8* KO cells could not. Western blot analysis showed markedly decreased levels of Ago2 in KO cells, making RNAi-based knockdown approaches inefficient (**Supp Fig 3A**). As an alternative, we used chemical inhibitors of the IFN pathway, acting upstream and downstream of IFN transcription. These included BX-795, which is a potent inhibitor of the TBK1 kinases that phosphorylates IRF3^59^, and the JAK1/2 inhibitor, ruxolitinib^60^. Both inhibitors resulted in decreased expression of ISGs in *DGCR8* KO cells, bringing their levels closer to those observed in WT cells (**Fig 2A, Supp Fig 3B**). These findings suggests that increased ISG production in *DGCR8* KO cells results in part from enhanced signalling upstream of IRF3 activation. In addition, we did not observe an upregulation in other components of the pathway upstream of IRF3/7, such as MAVS (**Supp Fig 3C**), suggesting that neither this component of the RLR pathway nor the general miRNA deficiency are the primary drivers for the activation of the IFN response in human *DGCR8* KO cells.

**FIGURE 3.**
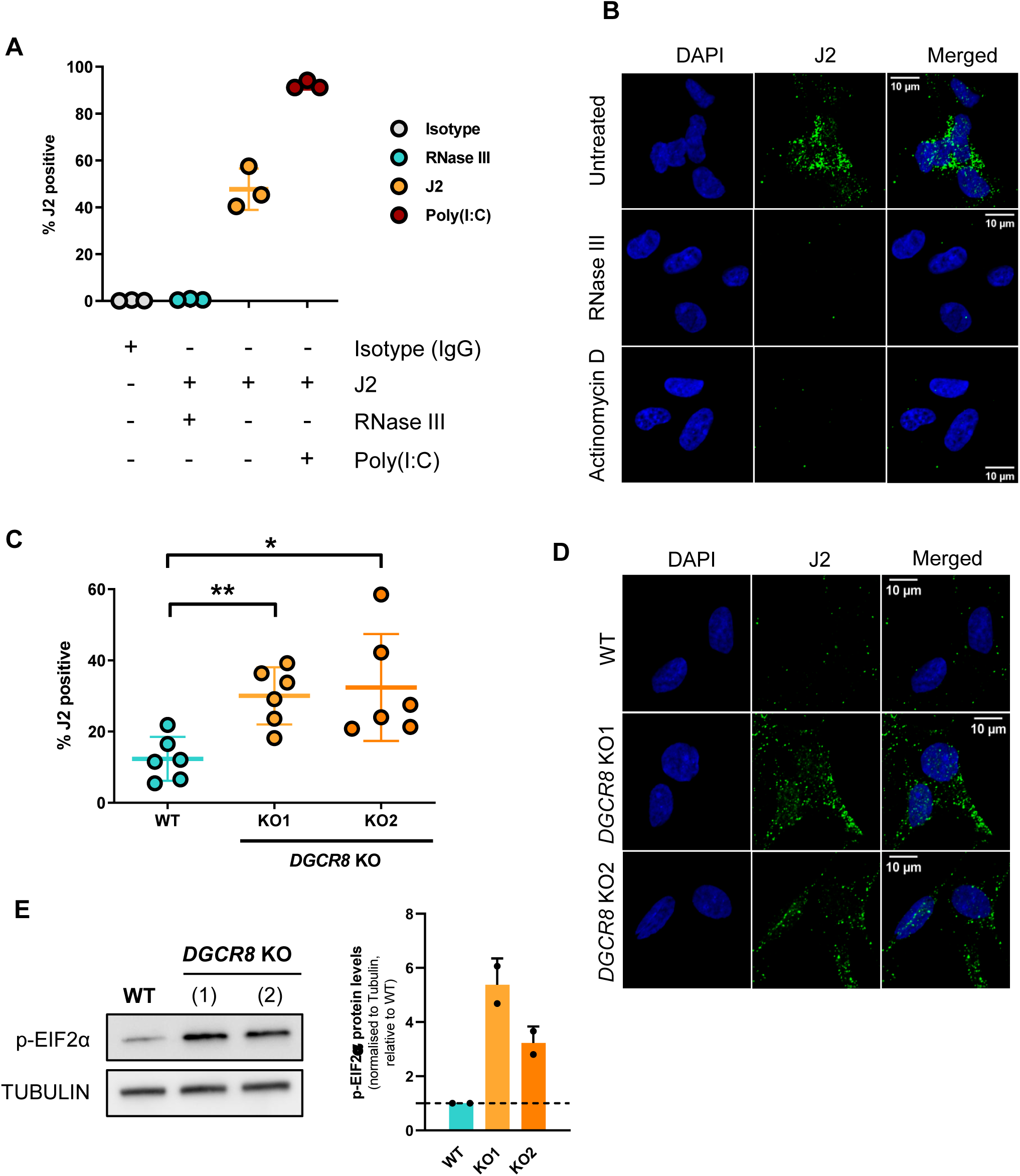
*DGCR8* KO cells accumulate dsRNAs in the cytoplasm. a. Flow cytometry quantification of J2 positive cells in PA-1 cells stained with J2 in basal conditions (yellow), in the presence of RNase III (blue) and after transfection with poly(I:C) (red), or Isotype IgG antibody control (grey). b. Representative immunofluorescence images of *DGCR8* KO1 cells after treatment with RNase III or actinomycin D followed by J2 staining. DAPI is used for nuclear counterstain. Scale bar represents 10 µm. c. Flow cytometry quantification of J2 positive cells in WT (blue) and two independent *DGCR8* KO clones in unstimulated conditions (yellow and orange). d. Representative immunofluorescence images of WT and two *DGCR8* KO clones stained with J2 antibody. DAPI is used for nuclear counterstain. Scale bar represents 10 µm. e. Western blot analyses of endogenous p-EIF2α in PA-1 WT and two *DGCR8* KO clones. Tubulin is used as loading control. On the right, quantification of p-EIF2α mark relative to Tubulin levels in two independent biological replicates. The dashed line represents the WT mean. Error bars plot mean ± SD of at least two replicates. *p < 0.05, **p < 0.01, ***p < 0.001, ****p < 0.0001. P-values are determined by one-way ANOVA followed by Dunnett multiple comparison post-hoc test.

To further investigate the origin of increased ISG expression, we compared the ability of WT and *DGCR8* KO cells to respond to immunogenic dsRNA, exogenous IFN, and DNA. Stimulation with poly(I:C), a dsRNA analogue typically recognised by the RLR melanoma differentiation-associated protein 5 (MDA5), resulted in increased activation of the IFN response in *DGCR8* KO cells (**Fig 2B**). Increased ISG expression in *DGCR8* KO cells was also observed upon exogenous IFN-β stimulation (**Fig 2C**). This upregulated responsiveness is consistent with the enhanced resistance to infection observed in *DGCR8* KO cells (**Fig. 1J**) and supports the notion that increased basal IFN levels in DGCR8-deficient cells prime them for an amplified immune response.

In contrast to this, the viral DNA mimic G3-YSD did not result in IFN activation in PA-1 cells, whereas it did in the control microglial cell line (BV-2), suggesting that PA-1 cells do not have a functional cGAS/STING signalling pathway (**Fig 2D**). In agreement with the lack of DNA sensing, we treated WT and *DGCR8* KO cells with a DNA damage agent, etoposide, since DNA damage has also been linked to IFN activation by several studies^61^. We found no increased ISGs expression after etoposide treatment, despite accumulation of DNA damage as demonstrated by gH2AX staining (**Fig 2E)**. To further confirm that DNA is not the trigger for the IFN response in *DGCR8* KO cells, we tested the effect of reverse transcriptase inhibitors. These inhibitors block cDNA synthesis derived from transposable elements and have been shown to prevent IFN activation in other contexts^62^. We blocked cDNA synthesis using two reverse transcriptase inhibitors lamivudine and emtricitabine, which are specific to LINE-1, the only active retrotransposon in humans that still encodes a functional reverse transcriptase domain^21^. None of these inhibitors reduced ISG expression in *DGCR8* KO cells (**Supp Fig 3D**). These results together suggest that increased expression of ISGs in *DGCR8* KO cells results from constitutive IFN production through the RLR signalling pathway, independent of DNA sensing.

### Constitutive IFN response is driven by dsRNA accumulation in *DGCR8* KO cells

Considering that the RLR pathway is constitutively activated in the absence of the dsRNA-binding protein DGCR8, we tested whether *DGCR8* KO cells were accumulating endogenous dsRNA. To this end, we developed several assays based on the J2 antibody, a monoclonal antibody that recognises dsRNA of at least 40 bp^63^. We first confirmed the specificity of the antibody for dsRNA quantification by flow-cytometry and immunofluorescence. Treatment with the dsRNA-specific endonuclease RNase III, which cleaves dsRNA resulting in fragments around 21-23 bp, prevents J2 staining (**Fig 3A-B**). In agreement with the ability of this antibody to recognise dsRNA, J2 staining by flow cytometry was increased after transfecting cells with the dsRNA poly(I:C) (**Fig 3A**). Using these approaches, we observed that *DGCR8* KO cells significantly accumulate more dsRNA than WT cells (**Fig 3C-D**). The majority of the dsRNA signal was enriched in the cytoplasm, where dsRNA sensing typically occurs (**Fig 3B, 3D**). Importantly, the dsRNA signal in *DGCR8* KO cells was lost after 2h of actinomycin D treatment, indicating that these dsRNAs arise from active transcription (**Fig 3B**). Consistent with the accumulation of cytoplasmic dsRNA, *DGCR8* KO cells also showed increased levels of phosphorylated eIF2α (p-eIF2α) (**Fig 3E**), which is phosphorylated by the dsRNA sensor protein kinase R (PKR) upon dsRNA recognition^64^. All these findings suggest that *DGCR8* KO cells accumulate dsRNA in the cytoplasm, and this can be sensed by several sensors, including the RLR pathway and PKR.

### 3’ UTR-localised retrotransposons form dsRNA in DGCR8 KO cells

To identify the endogenous cellular dsRNAs responsible for triggering the IFN response, we immunoprecipitated cytoplasmic dsRNAs using the J2 antibody followed by high-throughput sequencing (dsRNA-Seq). At a global level, the majority of reads mapped to protein-coding genes (84.3% in KO1, 85.1% in KO2), a smaller percentage of reads mapped to repetitive sequences (13.8% in KO1, 13% in KO2) and only ∼1% was assigned to other non-coding features (**Fig 4A**). Next, we performed differential enrichment analyses for both *DGCR8* KO dsRNA-seq relative to WT. While protein-coding genes were not generally more enriched in the dsRNA-seq of *DGCR8* KO cells (**Supp Fig4A-B** and **Supp Table S4-5**), individual TE loci were predominantly enriched in both *DGCR8* KO lines compared to WT (**Fig 4B-C** and **Supp Table S6-7**). Detailed analysis of the dsRNA-forming TE loci enriched in KO cells revealed that these primarily mapped to SINE (∼50%) and LINE (∼25%) retrotransposons (**Fig 4D**).

**FIGURE 4.**
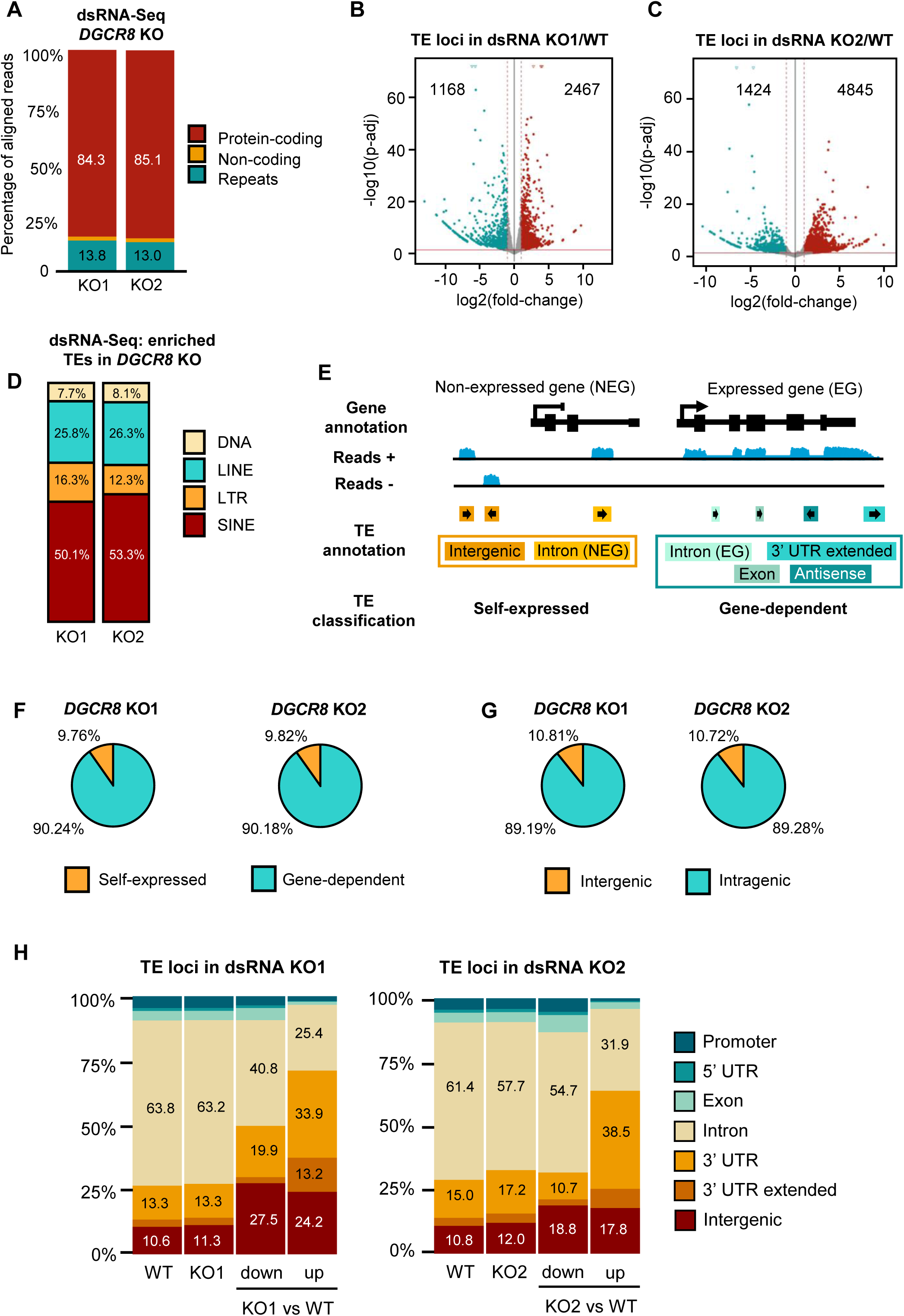
3’ UTR-localised transposable elements form dsRNA in *DGCR8* KO cells. a. Read distribution for the J2-based immunoprecipitation of dsRNA (dsRNA-seq) using two independent *DGCR8* KO clones. Reads mapping to more than one category are counted in each. b. Differential TE loci expression in dsRNA-seq IPs from *DGCR8* KO1 cells vs WT. Blue represents significantly downregulated TE loci (log2FC < −1 and p-adj < 0.05), red represents significantly upregulated (log2FC > 1 and p-adj < 0.05). c. Same as (b) for *DGCR8* KO2 dsRNA-seq. d. Fraction of enriched TE loci (log2FC > 1 and p-adj < 0.05) in *DGCR8* KO1 and KO2 dsRNA-Seq. e. Representation of TE classification; ‘self-expressed’ TEs are those annotated in intergenic regions or introns of a non-expressed protein-coding gene. Gene-dependent TEs are those inserted within exons, 5’ UTRs, 3’ UTRs, extended 3’ UTRs, introns of an expressed gene, or expressed on antisense orientation. f. Fraction of self-expressed vs gene-dependent dsRNA-forming TE loci enriched in both *DGCR8* KO clones. g. Fraction of intragenic vs intergenic dsRNA-forming TE loci enriched in both *DGCR8* KO clones. h. Genomic distribution of dsRNA-forming TE loci in WT, *DGCR8* KO1 and KO2 cells, versus those enriched (up) in KO1 or KO2 vs WT (log2FC > 1 and p-adj < 0.05) and depleted (down) in KO1 or KO2 vs WT (log2FC > −1 and p-adj < 0.05). Proportions represent the average of three biological replicates.

TEs are highly abundant throughout the human genome, particularly in intergenic regions and introns, but can also be found within coding sequences and UTRs^5^. As a result, a high proportion of the RNA found in cells that contain TE sequences originates from transcription within the gene where they are inserted and does not reflect transcription from the TE promoter itself^65^. This is of particular importance when studying SINEs, specially Alus, which are typically embedded within gene-transcripts^12^. Based on a previously described pipeline^37^, we established a method to determine if TE-derived dsRNA originated from TE transcripts arising from their own promoters (self-expressed) or alternatively, from TE sequences embedded within other transcripts (gene-dependent). We classified TE annotations based on their location relative to the genes’ position and reads’ orientation. TEs inserted within an expressed gene or transcribed in the antisense direction relative to their own sequence were considered gene-dependent. Conversely, TEs expressed in the sense orientation and located in intergenic regions or within introns of non-expressed genes were classified as self-expressed (**Fig 4E**). We found that most TEs in the dsRNA-Seq originated from protein-coding genes and were not self-expressed elements (∼90%, **Fig 4F**). Consistently, when individual TE copies were classified depending on their genomic location, we found that most immunoprecipitated dsRNA-forming TE loci were intragenic for both KO cell lines, mostly originating from the 3’ UTR of protein-coding genes (**Fig 4G-H**). Taken together, these findings suggest that SINE elements within the 3’ UTR of protein-coding genes are enriched in the dsRNA fraction of *DGCR8* KO cells, and could be responsible for triggering a constitutive IFN response.

### DGCR8 binds and regulates dsRNA-derived SINE-rich mRNAs

Considering the specific enrichment of SINE-containing 3’ UTR dsRNA in KO cells, we sought to determine whether this regulation could be mediated by direct binding of DGCR8 to these transcripts. We and others have previously demonstrated that DGCR8 not only binds pri-miRNAs but also other types of transcripts, including protein-coding RNAs^15,18,43,55,66,67^. In addition, DGCR8 binds TE-derived RNAs, particularly LINEs and the SINEs, mostly Alus^18^, indicating that DGCR8 could bind TE-sequences contained within mRNAs.

We first investigated if binding of DGCR8 to protein-coding genes was promoted by the presence of TE sequences. Using the HITS-CLIP dataset for human DGCR8^43^, we found that DGCR8-bound genes had a significantly higher number of TEs than those not bound (**Fig 5A**). To correct for potential bias caused by gene length, we calculated the density of TE-derived sequences per gene. Again, we observed that genes bound by DGCR8 had a significantly higher density of TEs than those not bound (**Fig 5B**). When discriminating by TE families, we found that both L1s (LINEs) and Alus (SINEs) were particularly enriched in genes bound by DGCR8, with significantly higher number of SINEs and LINEs located in the 3’ UTR of genes bound by DGCR8 compared to unbound genes. The number of LTRs remained similar in both subgroups (**Fig 5C** and **Supp Fig 4C-D**). These results indicate that DGCR8 binding to mRNAs is influenced by the presence of TEs, in particular SINE and LINE elements. To experimentally confirm this observation, we validated binding of DGCR8 to genes forming SINE-derived dsRNA from their 3’ UTRs. Here, we selected genes that contain multiple SINE elements, mostly Alu, in both sense and antisense orientation which can form dsRNA^11^. Immunoprecipitation of endogenous DGCR8 followed by RT-qPCR confirmed DGCR8 binding to these mRNAs (**Fig 5D**). These findings reveal that DGCR8 preferentially binds protein coding RNAs containing SINE elements, particularly in their 3’ UTR, suggesting a specific role for DGCR8 in the regulation of TE-enriched transcripts.

**FIGURE 5.**
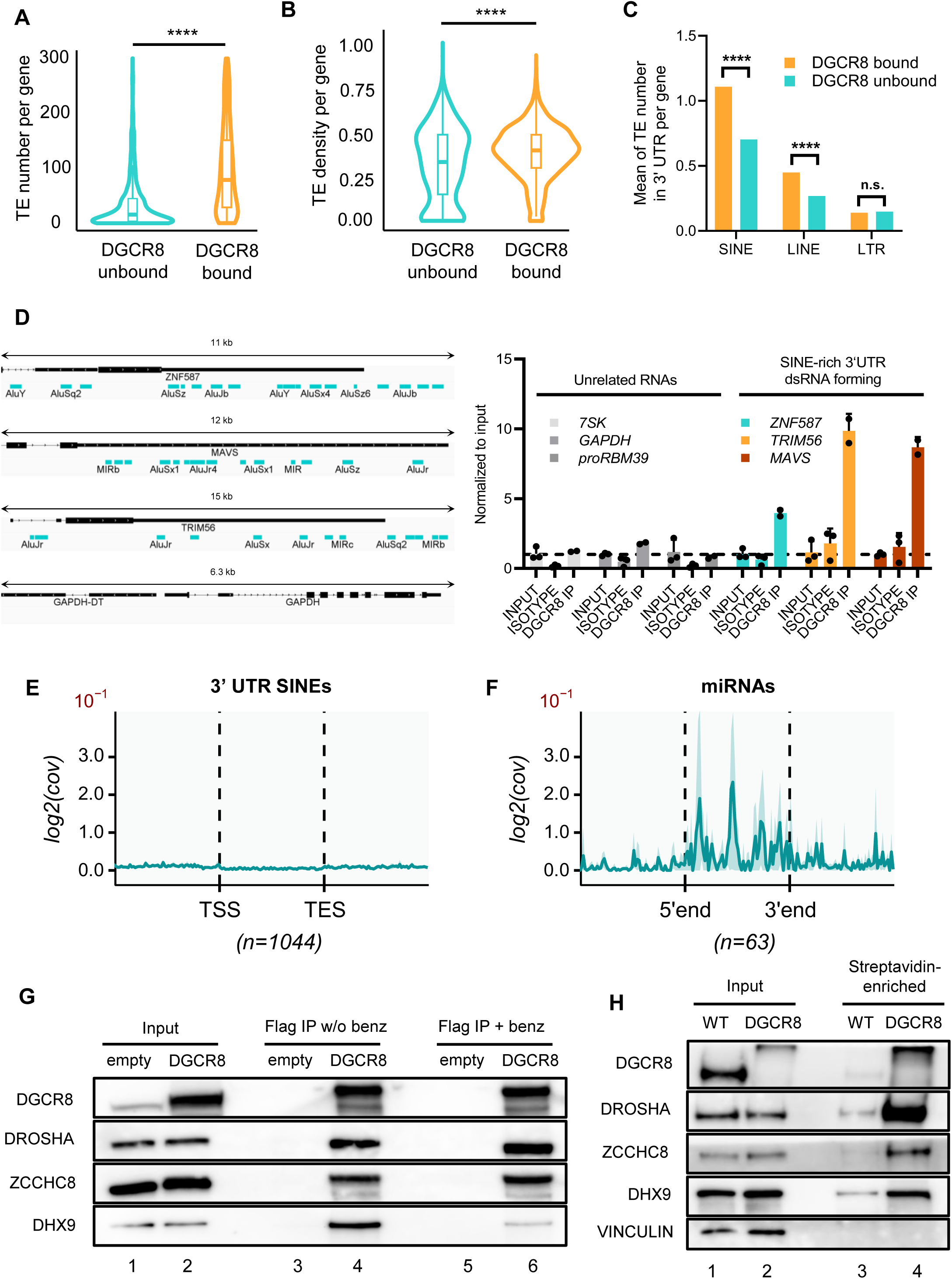
DGCR8 binds SINE-rich mRNAs. a. Absolute number of TEs per gene bound by DGCR8 and unbound defined by HITS-CLIP of DGCR8 (Macias et al 2012). b. Density of TEs per genes bound and unbound by DGCR8. c. Average number of SINE, LINE and LTR elements in the 3’ UTR of genes bound and unbound by DGCR8. d. On the left, genome browser view of selected 3’ UTRs of protein-coding genes. Annotated TEs are shown below in blue. On the right, RT-qPCR analysis of associated transcripts (*ZNF587, TRIM56, MAVS*) with immunoprecipitated endogenous DGCR8 (RIP). *RN7SK, GAPDH* and *RBM39* PROMPT serve as negative controls. Relative enrichment over input is represented (n=3). e. Aggregate plot of mean pA+/− RNA 3’ end-seq from HeLa cells (Gockert et al. 2022). Signal was plotted over 300 bp TSS-upstream to 300 bp TES-downstream regions of annotated SINE elements in 3’ UTRs of mRNAs with logFC >1 & padj <0.05 in *DGCR8* KO2/WT dsRNA-Seq (n=1044). The transcriptional unit (TU) body area (TSS to TES) was normalised to 300 bp for each SINE. Mean values and 90% confidence intervals (lighter shade around the curves) of log2-transformed coverage data are displayed in the aggregate plot with binning of 5 and a 10^−1^multiplication factor. f. Aggregate plot of mean pA+/− RNA 3’ end-seq from HeLa cells (Gockert et al. 2022). Signal was plotted over 25 bp upstream to 25 bp downstream regions of expressed mature miRNA (n=63). Mean values and 90% confidence intervals (lighter shade around the curves) of log2-transformed coverage data are displayed in the aggregate plot with binning of 5 and a 10^−1^multiplication factor. g. Immunoprecipitation of overexpressed FLAG-DGCR8 in HeLa cells and co-immunoprecipitated DROSHA, ZCCHC8 and DHX9 levels by western blot. Benzonase treatment serves to test RNA-dependent interactions. h. Proximity labelling assay of WT and mTurboID-DGCR8 mESCs. Biotinylated proteins were enriched using streptavidin beads, followed by western blot against DGCR8, DROSHA, ZCCHC8, DHX9 and vinculin as a control. In a-c, p-values are determined by Welch Two Sample t-test. *p < 0.05, **p < 0.01, ***p < 0.001, ****p < 0.0001.

We next asked whether the interplay between DGCR8 and TE-rich sequences and consequent dsRNA accumulation was mediated by the loss of Drosha. For this, we analysed 3’ end RNA-sequencing data looking for evidence of Drosha cleavage within SINE elements contained in the 3’ UTRs of cells. To do so, annotated SINE elements enriched in the dsRNA-Seq of *DGCR8* KO cells were selected as potential targets. No peaks indicative of endonucleolytic cleavage were detected within SINE sequences in 3’ UTRs (**Fig 5E),** while signs of cleavage in primary miRNAs, the canonical Drosha substrates, were observed (**Fig 5F**). These findings suggest that Drosha is not involved in cleaving dsRNA structures within 3’ UTRs.

Given the absence of cleavage in SINE-enriched dsRNAs, we explored whether alternative mechanisms might regulate the extent of dsRNA formation from these sequences. Several studies have reported the protein interactome of human DGCR8 by immunoprecipitation followed by mass spectrometry^15,67^. The RNA helicase DHX9 has been found as a DGCR8 interactor in two independent mass-spectrometry experiments^15,67^. This helicase has also been shown to resolve dsRNA structures from Alu elements in 3’ UTRs^68^. To test if DHX9 could cooperate with DGCR8, we performed co-immunoprecipitation (IP) analyses. We confirmed that DGCR8 binds DHX9 by co-IP using HeLa cell extracts overexpressing FLAG-DGCR8. This interaction was partially sensitive to the activity of benzonase, indicating that co-IP of DGCR8 with DHX9 was RNA-dependent and suggesting that both proteins bind the same RNAs (**Fig 5G**). To validate the DGCR8-DHX9 interaction in living cells using orthologous methods, we performed a proximity labelling assay in mESCs using the mTurboID system^69^. A 2xHA-tagged miniTurbo (HA-mTurbo) biotin ligase enzyme was genomically integrated to the N-terminus of the endogenous DGCR8 gene in mESCs using CRISPR/Cas9. Following the addition of biotin, biotinylated proteins proximal to mTurbo-DGCR8 were isolated using streptavidin affinity purification. Western blot analysis detected DHX9 along with other well-known DGCR8 interactors, including Drosha and ZCCHC8^70^, thereby confirming that DGCR8 and DHX9 are in close physical proximity in living cells (**Fig 5H**).

Together, this led us to hypothesise that DGCR8 binding to SINE-rich genes impacts dsRNA formation independently of Drosha cleavage and potentially by cooperation with the RNA helicase DHX9. In the absence of DGCR8, dsRNA-forming TE-rich mRNAs accumulate in the cytoplasm triggering dsRNA sensing by RLR and PKR.

### Loss of DGCR8 leads to immune system dysregulation in 22q deletion syndrome cells

Mutations in dsRNA binding proteins, such as MDA5 or ADAR1, lead to hyperactivation of the IFN response, resulting in autoimmune and autoinflammatory diseases^62^. Considering this link and the role of DGCR8 in suppressing the IFN response in human cells, we next investigated if diseases characterised by altered DGCR8 levels could also display aberrant IFN responses. We focused on the 22q11.2 deletion syndrome (22qDS), a genetic syndrome where DGCR8 expression levels are affected. The disease is caused by a hemizygous deletion in chromosome 22 which encompasses several genes, including *DGCR8*. As a consequence, 22qDS patients only harbour a single functional *DGCR8* allele, and we have recently confirmed that losing a single *DGCR8* copy is enough to alter function in human embryonic stem cell models^20^. To test the importance of DGCR8 levels in controlling the IFN response, we first assessed the behaviour of heterozygous *DGCR8* (HET) PA-1 cells in comparison to WT and *DGCR8* KO cells. When cells can only produce DGCR8 protein from a single allele, they display an intermediate phenotype of IFN activation when challenged with the dsRNA poly(I:C) (**Fig 6A**). *DGCR8* HET cells also display an intermediate phenotype in the susceptibility to RNA viruses, such as Influenza A Virus (IAV) (**Fig 6B**). These findings suggest that DGCR8-mediated control of the IFN response is dependent on *DGCR8* gene dosage.

**FIGURE 6.**
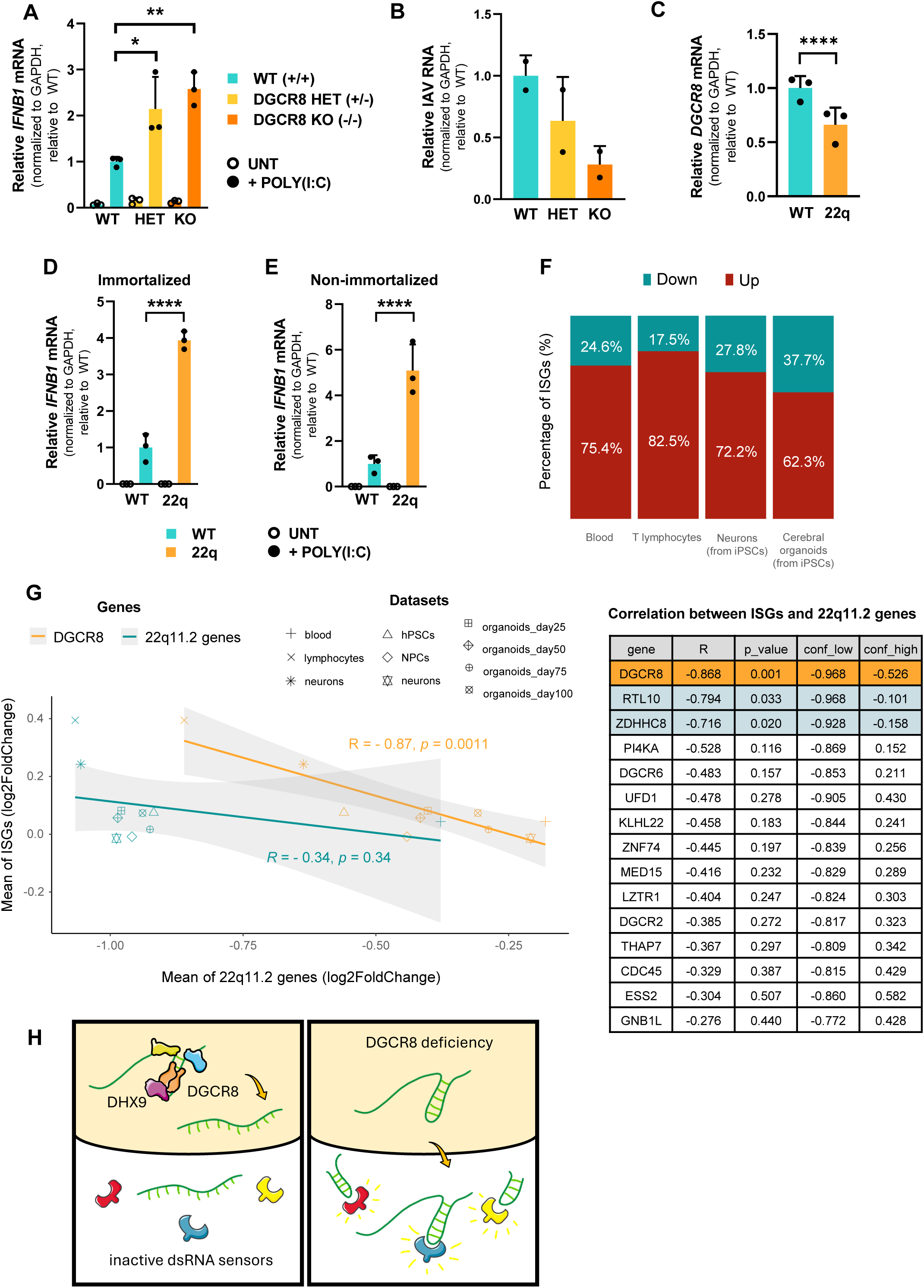
Loss of DGCR8 leads to immune system activation in 22qDS cells. a. Quantification of *IFNB1* mRNA levels by RT-qPCR in WT, *DGCR8* HET and *DGCR8* KO PA-1 after stimulation with poly(I:C) vs untreated. Data was normalised to *GAPDH* and relative to stimulated WT cells. b. Influenza A Virus RNA levels by RT-qPCR after infection of WT, *DGCR8* HET and *DGCR8* KO. *18S rRNA* was used as normaliser, data represented relative to WT cells. c. *DGCR8* mRNA levels by RT-qPCR in control (healthy cells) and 22qDS immortalised fibroblasts (GM07215F). Data are normalised to *GAPDH* and relative to WT cells. d. *IFNB1* expression levels by RT-qPCR in control (healthy cells) vs 22qDS immortalised fibroblasts (GM07215F), after poly(I:C) stimulation. Data are normalised to *RN7SK* and represented relative to stimulated healthy (WT) fibroblasts. e. *IFNB1* expression levels by RT-qPCR in control (healthy cells) vs 22qDS derived primary fibroblasts (GM02944), after poly(I:C) stimulation. Data are normalised to *GAPDH* and represented relative to healthy stimulated cells. f. Percentage of ISGs with a log2FC (22qDS/control) > 0 (up, red) or log2FC(22qDS/control) < 0 (down, blue) from the following datasets: blood (Lin et al. 2021), T-lymphocytes (Raje et al. 2022), iPSCs-derived neurons (Lin et al. 2016) and cerebral cortex organoids from 22qDS-iPSCs (Khan et al. 2020). g. Correlation between the log2FC (22qDS/control) of 22q11.2 deleted genes and ISGs. The log2FC of the mean of ISGs is plotted on the y-axis. The log2FC of the mean of 22q11.2 genes (blue) or *DGCR8* gene (orange) are plotted on the x-axis. Each dot represents one dataset, these being: blood (n=79 patients, n=68 controls) (Lin et al. 2021), T-lymphocytes (n=13 patients, n=6 controls) (Raje et al. 2022), iPSCs-derived neurons (n=8 patients, n=7 controls) (Lin et al. 2016), hPSCs, NPCs and neurons (n=20 patients, n=29 controls) (Nehme et al. 2022) and cerebral cortex organoids differentiated from 22qDS-iPSCs after 25, 50, 50 and 100 days (n=15 patients, n=14 controls) (Khan et al. 2020). On the right, a table representing the top15 22q11.2 genes high the highest correlation index (all genes in Supplementary Table S8). h. Proposed model: In healthy conditions (left), DGCR8 and dsRNA-BPs such as DHX9 bind and resolve dsRNA structures of TE-rich mRNAs. After unwinding, RNAs are exported to the cytoplasm, where cannot be recognised by RIG-I-like receptors. In the absence of DGCR8 (right), dsRNAs remain unresolved. In the cytoplasm, these dsRNA-rich mRNAs are recognised by RIG-I-like sensors triggering IFN activation. In RT-qPCR, bars represent the average of three biological replicates. Error bars plot mean ± SD. *p < 0.05, **p < 0.01, ***p < 0.001, ****p < 0.0001. P-values are determined by one-way ANOVA followed by Dunnett multiple comparison post-hoc test.

To test the relevance of the DGCR8-IFN axis in the disease context, we compared the status of the IFN response in immortalised primary fibroblasts from a 22qDS patient (GM07215F) and a healthy control. First, we confirmed that 22qDS-derived fibroblasts expressed reduced *DGCR8* mRNA levels (**Fig 6C**). Considering that basal levels of *IFNB1* are difficult to detect in fibroblasts, we challenged these patient-derived cell lines with exogenous dsRNA. This resulted in significantly increased *IFNB1* levels in 22qDS fibroblasts compared to healthy controls following stimulation (**Fig 6D**). To ensure that these differences were not biased by individual-specific effects or immortalisation, we also challenged primary, non-immortalised fibroblasts derived from a different patient (GM02944). Again, we found higher expression of *IFNB1* in 22qDS human primary fibroblasts upon dsRNA stimulation compared to control (**Fig 6E**).

To extend these findings using a larger cohort of patients, we analysed the IFN signature in several 22qDS-patient datasets. We first analysed the microarray-based gene expression data of peripheral blood from 79 patients and 68 age and sex-matched controls^50^ and the RNA sequencing datasets on T-lymphocytes of patients (n= 13) and healthy controls subjects (n=6)^47^ and we found a global upregulation of ISGs compared to control cells (**Fig 6F, Supp Fig 5A**). Next, we performed Gene Set Enrichment Analysis (GSEA) of the differentially expressed genes in 22qDS patients and found that all ontologies enriched in the upregulated genes belonged to ‘immune system activation’ (**Supp Fig 6**), indicating that 22qDS blood cell types exhibit an exacerbated immune response.

Given the link between 22qDS and neuronal disorders^49,71,72^ and the reduced tolerance of neurons to dsRNA accumulation^73^, we also investigated the presence of ISG signatures in neuronal tissues derived from 22qDS patients, including cerebral cortex organoids generated from 22qDS-iPSCs and iPSCs-derived neurons, using available RNA-seq datasets^46,49^. Similar to blood cell types, both neural models showed a global upregulation of ISGs (**Fig 6F, Supp Fig 5B**). Again, all ontologies enriched in the upregulated genes in 22qDS neurons belonged to immune system activation (**Supp Fig 7A**). Also, STAT2 target gene ontology was the most enriched transcription factor network in upregulated genes (**Supp Fig 7B**) indicating an activation of the JAK/STAT pathway in neurons derived from 22qDS cells.

22qDS involves the deficiency of up to 50 genes^74^, however, not all genes contribute equally to the penetrance of the syndrome. *DGCR8* is one out of the five genes affected by the deletion that is also predicted to be haploinsufficient in humans^20^. For critical genes in heterozygosity, cells often undergo selection for those expressing the remaining functional allele at higher levels^75^. Thus, we studied the depletion of 22qDS genes in the four studied datasets and found DGCR8 among the least depleted 22qDS genes (**Supp Fig 8A**). Interestingly, we observed that datasets with a higher ISG upregulation correlated with lower *DGCR8* expression levels (**Supp Fig 8B**). To increase robustness, we included six more datasets of 22qDS-derived cell types in this analysis. This confirmed that *DGCR8* expression was inversely correlated with the expression of ISGs (**Fig 6G**). We studied the correlation of ISGs expression with each 22qDS gene and found that among the 31 expressed 22qDS genes, only three showed a significant negative correlation with ISGs, with *DGCR8* being the gene with the highest and most significant correlation (R = - 0.868, p = 0.001) (**Fig 6G** and **Supp Table S8**).

These results together indicate that cells derived from 22qDS patients have a basal upregulation of the IFN pathway, and that alterations in *DGCR8* levels seem to be the main contributor to ISG upregulation in 22q11.2DS patients.

## Discussion

Mammalian cells must carefully regulate the production and recognition of endogenous dsRNA to avoid activating antiviral defence pathways in the absence of an actual pathogen. One of the primary responses to dsRNA is the production of IFNs, which can be harmful if inappropriately activated, as in the case of type I interferonopathies^76^. Beyond the IFN response, dsRNA recognition can trigger several other potent cellular responses, including the activation of integrated stress response (ISR). In this pathway, the dsRNA sensor PKR binds dsRNA and subsequently phosphorylates the initiation factor of translation eIF2α^77,78^. Our data suggest that at least both the IFN and the ISR response become activated upon DGCR8 loss, consistent with the cytoplasmic dsRNA accumulation observed in *DGCR8* KO cells.

Our findings contribute to the growing field of dsRNA binding proteins whose malfunctioning can cause human disease. For instance, MDA5, RIG-I and ADAR1 are mutated in Aicardi-Goutières syndrome causing hyperactivation of the IFN response^79^. Previous studies had shown that DGCR8 is implicated in suppressing the IFN response in mouse embryonic stem cells (mESCS) and mouse macrophages^56,80,81^. However, whether this regulatory function operates in human models, contributes to disease, and what drives IFN activation in such contexts remained unexplored.

In this current study, we have extended our investigation to 22qDS, a condition associated with reduced DGCR8 expression. Immunological abnormalities of 22qDS patients, which include immunodeficiencies and increased risk of autoimmunity, have traditionally been attributed to alterations in the thymus^74^. Our results lead us to hypothesise that some of the immune alterations could also be caused by increased IFN production. Hyperactivation of the IFN response in these patients may exhibit incomplete penetrance and could vary depending on the levels of DGCR8 in each patient. Moreover, uncontrolled IFN production could also contribute to other clinical features observed in these patients, particularly those involving dysfunction of the central nervous system. Approximately 25% of the individuals with 22qDS develop schizophrenia, representing a 25-fold increased risk compared to the general population^74^. The association between neuroinflammation and schizophrenia is becoming increasingly recognised^82–84^. Supporting this hypothesis, neuroinflammation has been reported in blood-brain barrier models derived from 22qDS iPSCs^72^, yet the precise molecular mechanisms remain unclear. We found that neurons derived from 22qDS patients showed signs of IFN activation. Interestingly, neurons are a cell type particularly vulnerable to dsRNA accumulation, as they present higher load of endogenous dsRNA and, as a consequence, a lower threshold for IFN activation^73^. This sensitivity could render neurons especially susceptible to the depletion of dsRNA-binding proteins (dsRBPs), such as DGCR8. Significantly, among the approximately 50 genes deleted in 22qDS, *DGCR8* exhibits the strongest inverse correlation with the expression of ISGs. Further studies are needed to explore whether DGCR8 could serve as a prognostic marker for immune activation in 22qDS patients or non-22qDS schizophrenia patients, and potentially help designing a stratification framework where patients with lower DGCR8 expression levels could be intervened with immunomodulatory treatments.

Our results also expand our knowledge on the non-canonical roles of DGCR8, beyond miRNA biogenesis^15–18,67,85^. DGCR8 has been shown to repress TE expression by promoting heterochromatin formation^17^. At the post-transcriptional level, we previously showed that DGCR8 repressed retrotransposon mobilisation by binding TE-derived RNAs and inducing cleavage by Drosha^18^. Here, we demonstrate that the regulation of TE by DGCR8 also applies to TE copies inserted within genes. Although a few studies suggest that the Microprocessor complex can cleave some mRNAs, including DGCR8 mRNA itself^43,55,66^, here we propose that DGCR8 regulation of RNAs can occur without cleavage and involves interaction with other co-factors. In support of our findings, DGCR8 has recently been shown to recognise stem-loop structures within nascent mRNAs to sequester transcriptional coactivators^67^. Further supporting our hypothesis, DHX9 depletion was shown to cause accumulation of cytoplasmic dsRNA and triggering of the IFN response, consistently with DGCR8 and DHX9 cooperating to bind and resolve dsRNA structures^86^.

Although many studies have pointed out TEs as the main source of endogenous immunostimulatory dsRNA^62^, it is often not thoroughly assessed whether these RNAs originate from autonomous TE promoters or are embedded within transcripts derived from host genes. Here we have performed a detailed analysis to distinguish if TE-dsRNAs originate from self-expressed TEs or from those inserted within genes. For this, we have modified a published pipeline developed for zebrafish^37^ and, as a key modification, we have considered antisense TE reads as gene-dependent TEs. This distinction is critical when quantifying TE expression to avoid misinterpretation^65^, especially when analysing their potential to form dsRNA. In the case of DGCR8, most of the dsRNA-forming TEs originated from protein-coding transcripts and were enriched in Alu elements located in the 3’ UTRs. In agreement with the role of these element in triggering the IFN response, inverted-repeat Alus were also found to be the main MDA5 targets using RNase protection assays^11^. Furthermore, Alu elements seem to be the most abundant target of ADAR1, which is responsible of editing dsRNA structures to prevent their recognition by MDA5^10^.

To conclude, our results support a model by which DGCR8 binds dsRNA structures formed by TEs inserted in the 3’ UTR of mRNAs and contributes to resolving them in cooperation with other dsRBPs, such as DHX9 (**Fig 6H**). Our findings emphasize how a fine balance between dsRNA levels and dsRBP expression and function is crucial to preserve cellular homeostasis^87^. Under normal conditions, cellular dsRBPs effectively bind dsRNAs, keeping dsRNA sensors in an inactive state. However, an increase in dsRNA concentration, following infections or alterations in RNA metabolism, or alternatively, a deficiency in a dsRBP such as DGCR8, can disrupt this balance. As a result, dsRNAs are recognised by cytoplasmic dsRNA sensors, triggering a complex network of cellular responses. We propose that disruption of this balance and consequent IFN activation, is a critical factor to consider in conditions where DGCR8 function is compromised, including disorders such as 22qDS.

## Acknowledgements/Fundings

We thank Irene Muñoz-Blat for helping with initial characterization of *DGCR8* KO cells, and 22q11.2 patients associations in Andalucia and Levante for their support and trust. This work has been funded by the Wellcome Trust [221737/Z/20/Z and 107665/Z/15/Z]; the Royal Society grant [RGS\R1\191368] and The Wellcome Trust iTPA [PIII021] (to S.M.); Grant CNS2023-145402 funded by MICIU/AEI/10.13039/501100011033 and by “European Union NextGenerationEU/PRTR”; The Spanish ministry of science and innovation (PID2020-115033RB-I00); Career Integration Grant-Marie Curie (FP7-PEOPLE-2011-CIG-303812); the Andalusian regional government (PY20_00619 y A-CTS-28_UGR20) grants and donation to “Aula de estudios 22qDS” (to S.R.H.); Ministerio de Ciencia e Innovación, Agencia Estatal de Investigación and European Union [PRE2021-098878] and EMBO Scientific Exchange Grant [SEG11137] (to A.G.-G.); MRC—Precision Medicine fellowship (to L.K.); Darwin Trust fellowship (to P.C.). Work in the T.H.J. laboratory was supported by the Independent Research Fund Denmark - Medical Sciences and the Novo Nordisk Foundation (NNF, ExoAdapt Grant 31199).

## Data Accessibility

All high-throughput RNA sequencing datasets have been deposited in the GEO database. Both total and small-RNA seq from WT and *DGCR8* KO cells can be accessed at GSE294708. The dsRNA-seq data for both WT and *DGCR8* KO cells can be accessed GSE294748.

## Author contributions

S.M., S.R.H., A.G.-G. conceived the project and wrote the manuscript with help and feedback from all authors. A.G.-G. and P.C. performed most experiments, with the help of K.G., J.W., L.L.-O., P.T.-R and W.G.. Most bioinformatic data curation and raw analysis was performed by G.P. with the help of A.G.-G., A.I. and L.K.; P.G.M. performed the 22qDS datasets analyses. J.R. performed the 3’ end RNA-sequencing analyses. Resources were provided by S.M., S.R.H., and T.H.J.. This article is part of the doctoral thesis of A.G.-G. at the University of Granada (Spain).

---

**SUPPLEMENTARY FIGURE 1, related to Figure 1.**
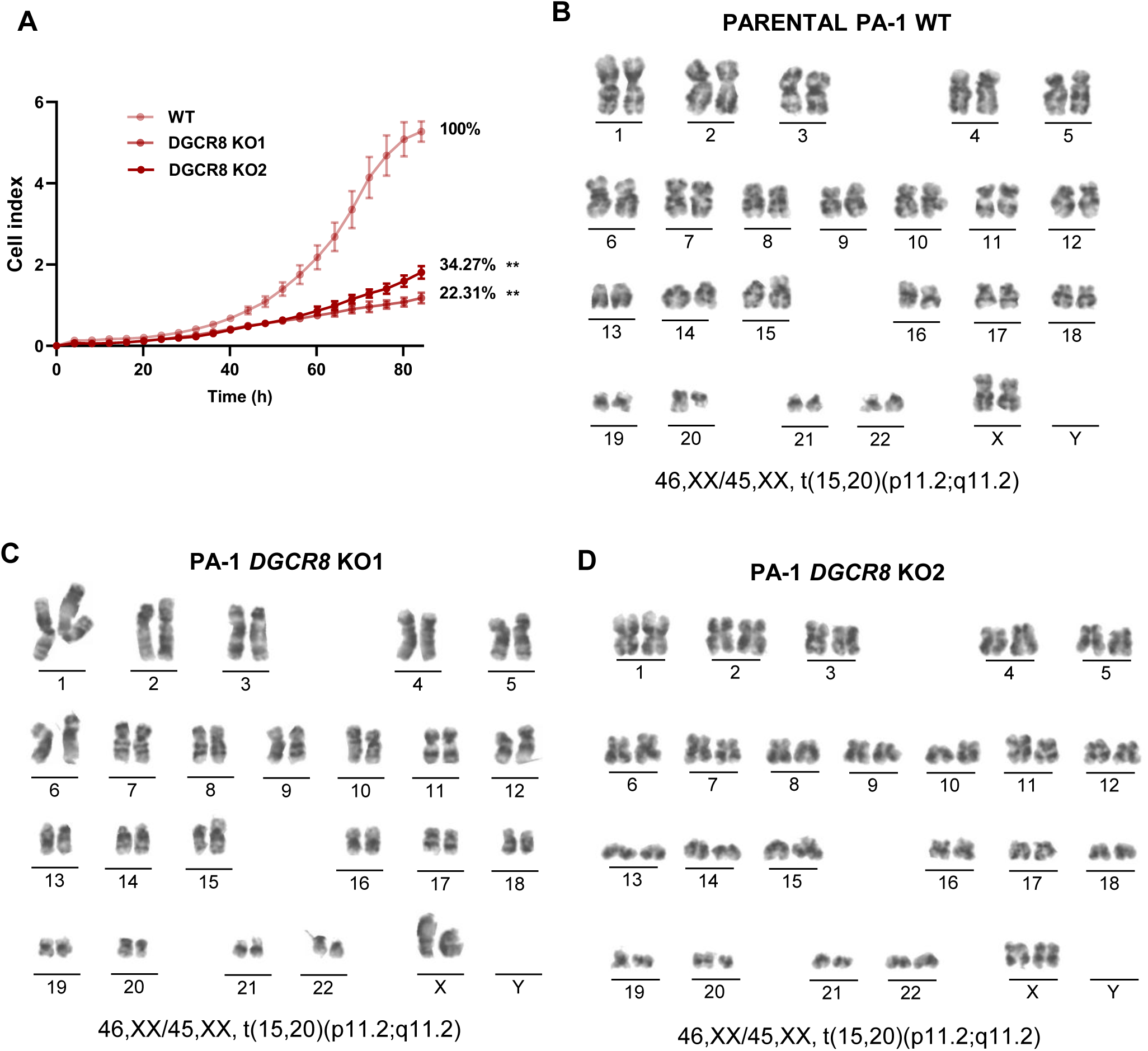
a. Raw cell index representing cell growth of PA-1 WT and two *DGCR8* KO clones for 84 hours. Error bars plot mean ± SD of three independent biological replicates. Percentage is calculated considering WT as 100% growth at final time. *p < 0.05, **p < 0.01, ***p < 0.001, ****p < 0.0001. P-values are determined by one-way ANOVA followed by Dunnett multiple comparison post-hoc test. b. Karyotype of human embryonic teratocarcinoma PA-1 WT cells obtained from metaphase chromosomes. c. Karyotype of human embryonic teratocarcinoma PA-1 *DGCR8* KO1 cells obtained from metaphase chromosomes. d. Karyotype of human embryonic teratocarcinoma PA-1 *DGCR8* KO2 cells obtained from metaphase chromosomes.

**SUPPLEMENTARY FIGURE 2, related to Figure 1.**
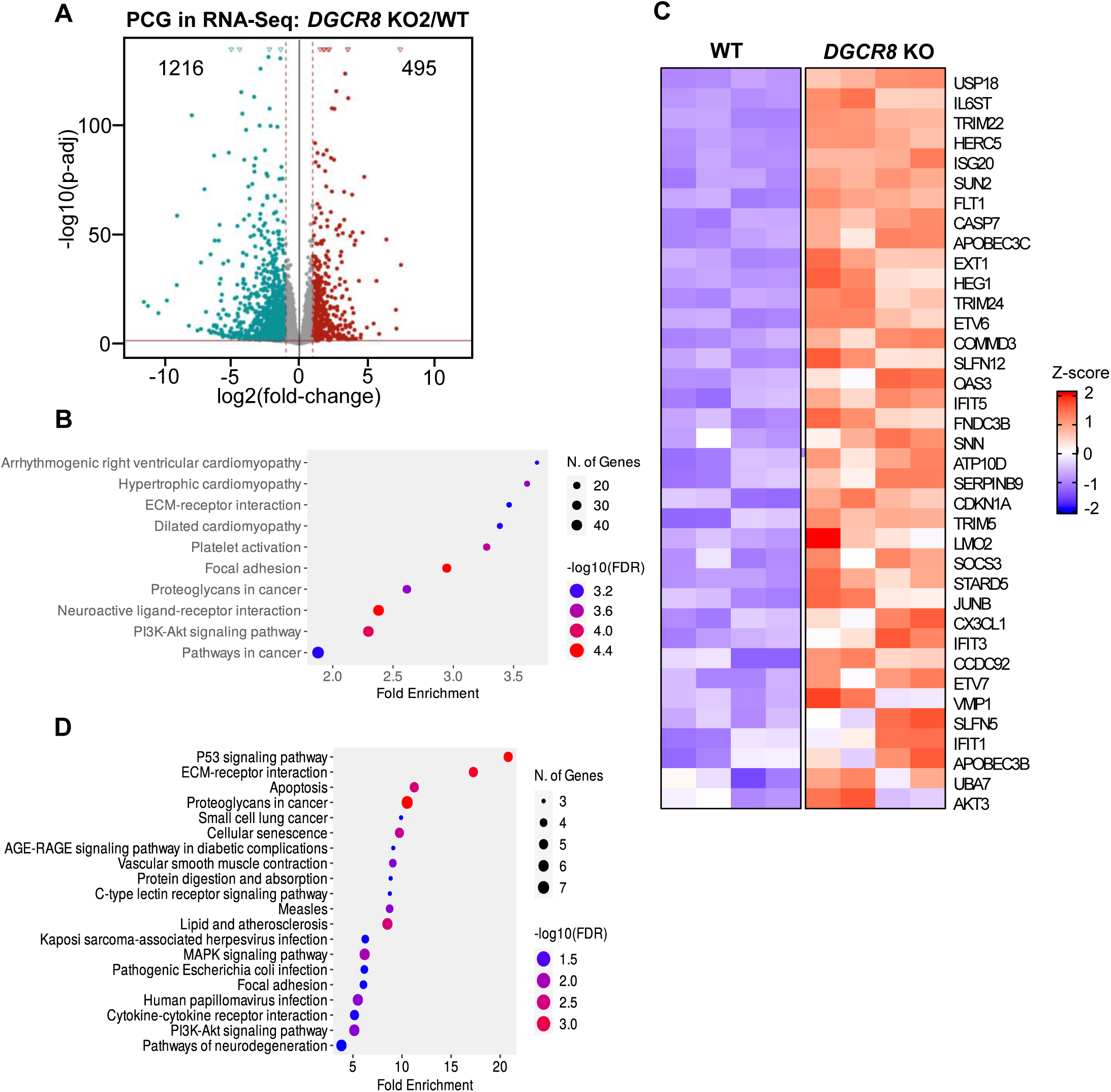
a. Volcano plot for differentially expressed (DE) protein-coding genes (PCGs) in *DGCR8* KO2 vs WT cells using total RNA-seq. Blue represents significantly downregulated PCGs (log2FC < −1 and p-adj < 0.05), red represents significantly upregulated (log2FC > 1 and p-adj < 0.05). b. KEGG enrichment analyses of downregulated genes in total RNA-Seq (log2FC < −1 and p-adj < 0.05) in *DGCR8* KO2 vs WT. c. Heatmap of upregulated (log2FC > 1 and p-adj < 0.05) ISGs in *DGCR8* KO2 cells by RNA-sequencing. Red indicates upregulation. d. KEGG enrichment analysis of upregulated proteins in *DGCR8* KO2 total cell proteomic extracts vs WT (log2FC > 1 and p-adj < 0.05).

**SUPPLEMENTARY FIGURE 3, related to Figure 2-3.**
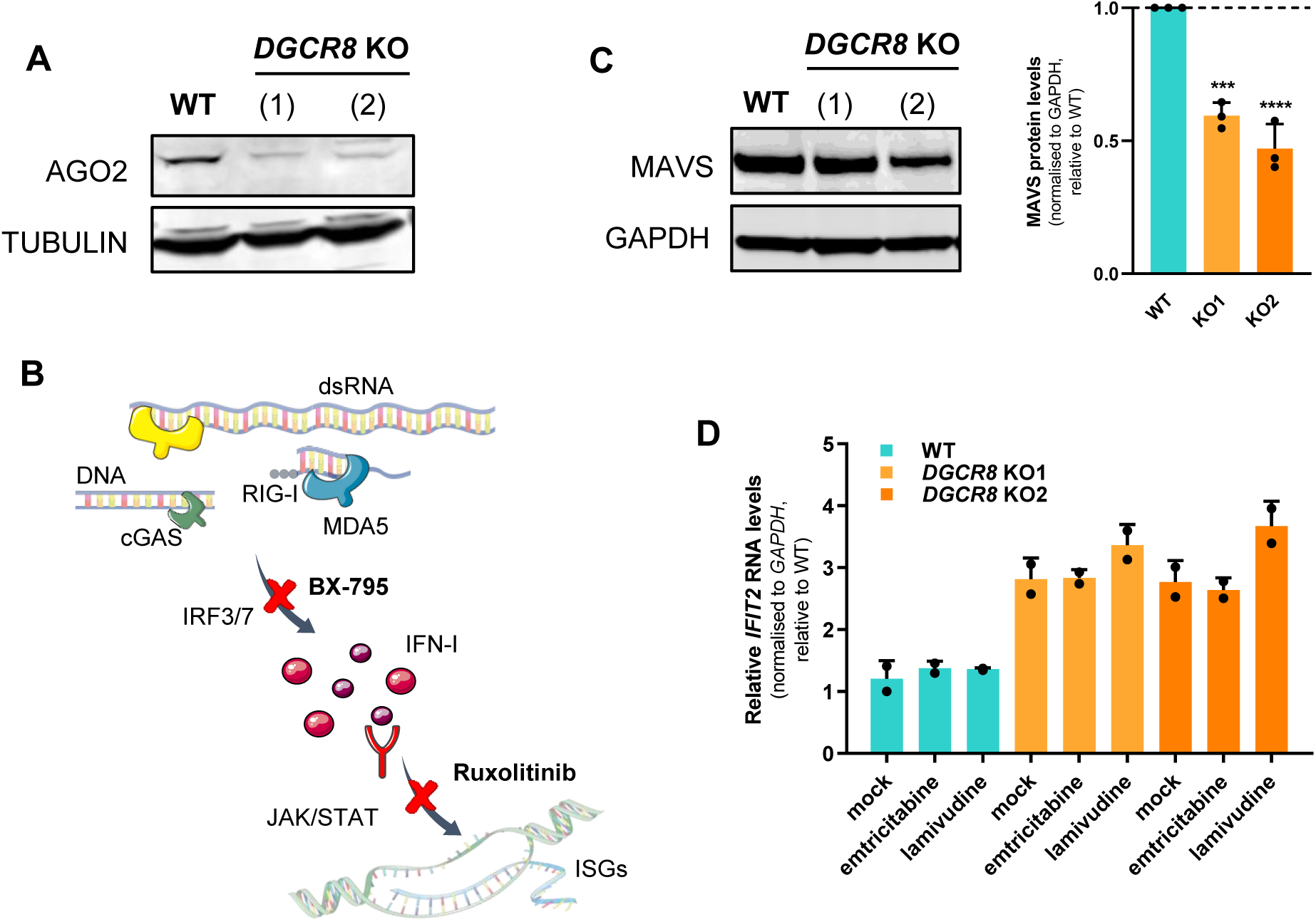
a. Western blot analyses of Ago2 protein levels in PA-1 WT and two independent *DGCR8* KO clones. Tubulin serves as a loading control. b. Scheme representing the pathways targeted by chemical inhibitors used in Figure 2A. c. Western blot analyses of MAVS protein levels in PA-1 WT and two independent *DGCR8* KO clones. GAPDH serves as a loading control (left). Quantification of MAVS protein levels using three independent biological replicates (right). Error bars plot mean ± SD. *p < 0.05, **p < 0.01, ***p < 0.001, ****p < 0.0001. P-values are determined by one-sample t-test. The dashed line represents the WT mean. d. *IFIT2* expression levels in WT and two *DGCR8* KO clones by RT-qPCR, after treatment with lamivudine, emtricitabine or vehicle. Data are normalised to *GAPDH* and expressed relative to WT mock sample. Error bars plot mean ± SD of two biological replicates.

**SUPPLEMENTARY FIGURE 4, related to Figure 4 and 5.**
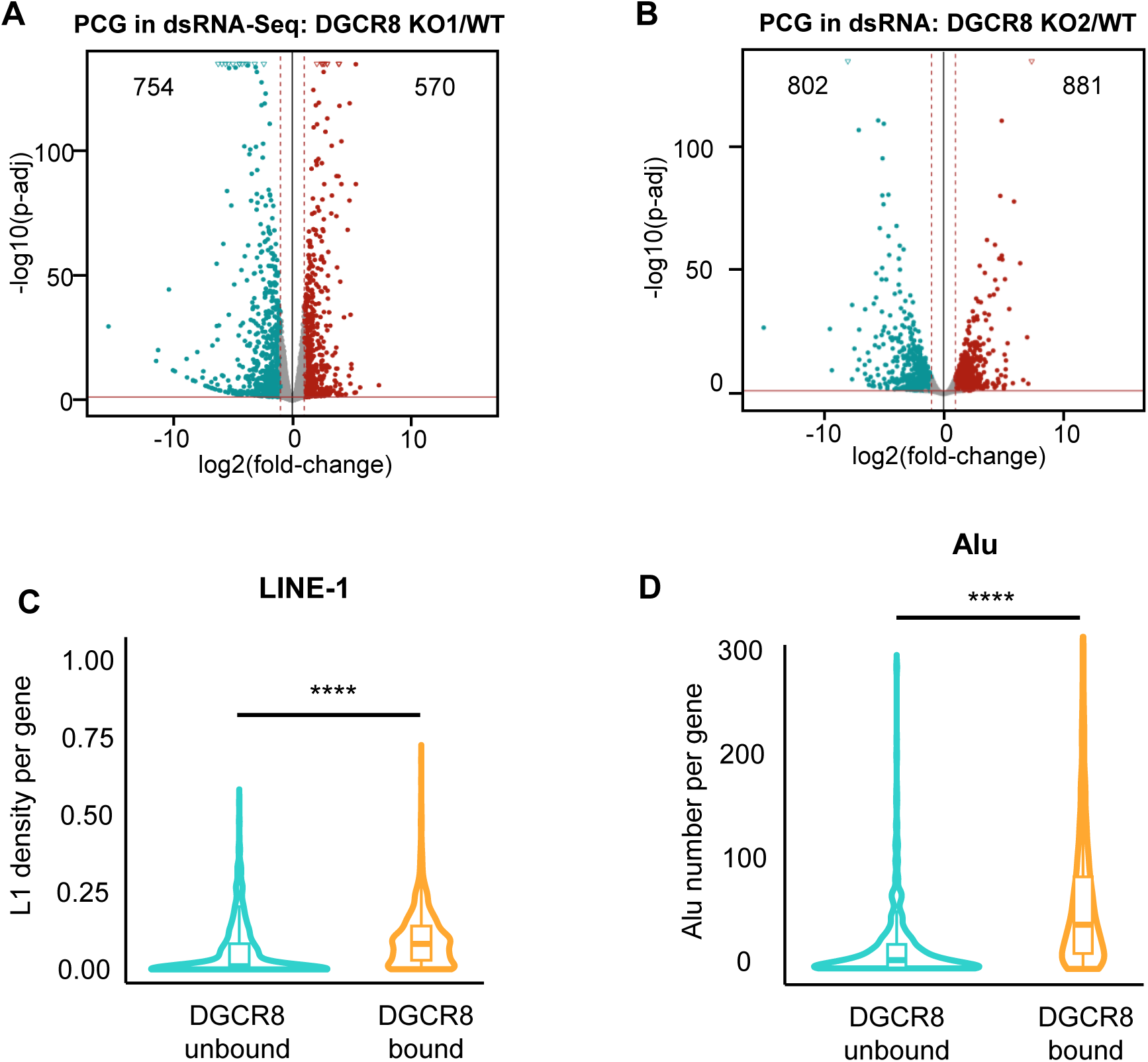
a. Differential expression analyses of protein-coding genes (PCGs) in dsRNA-seq of *DGCR8* KO1 vs WT cells. Blue represents significantly downregulated PCGs (log2FC < - 1 and p-adj < 0.05), red represents significantly upregulated (log2FC > 1 and p-adj < 0.05). b. Same as (a) for *DGCR8* KO2 vs WT cells. c. Violin plot representing the density of LINE-1 (L1) elements in DGCR8-bound vs non-bound genes defined by HITS-CLIP of DGCR8 from Macias et al. 2012. d. Violin plot representing the number of Alu elements in DGCR8-bound vs non-bound genes defined by HITS-CLIP of DGCR8 from Macias et al. 2012.

**SUPPLEMENTARY FIGURE 5, related to Figure 6.**
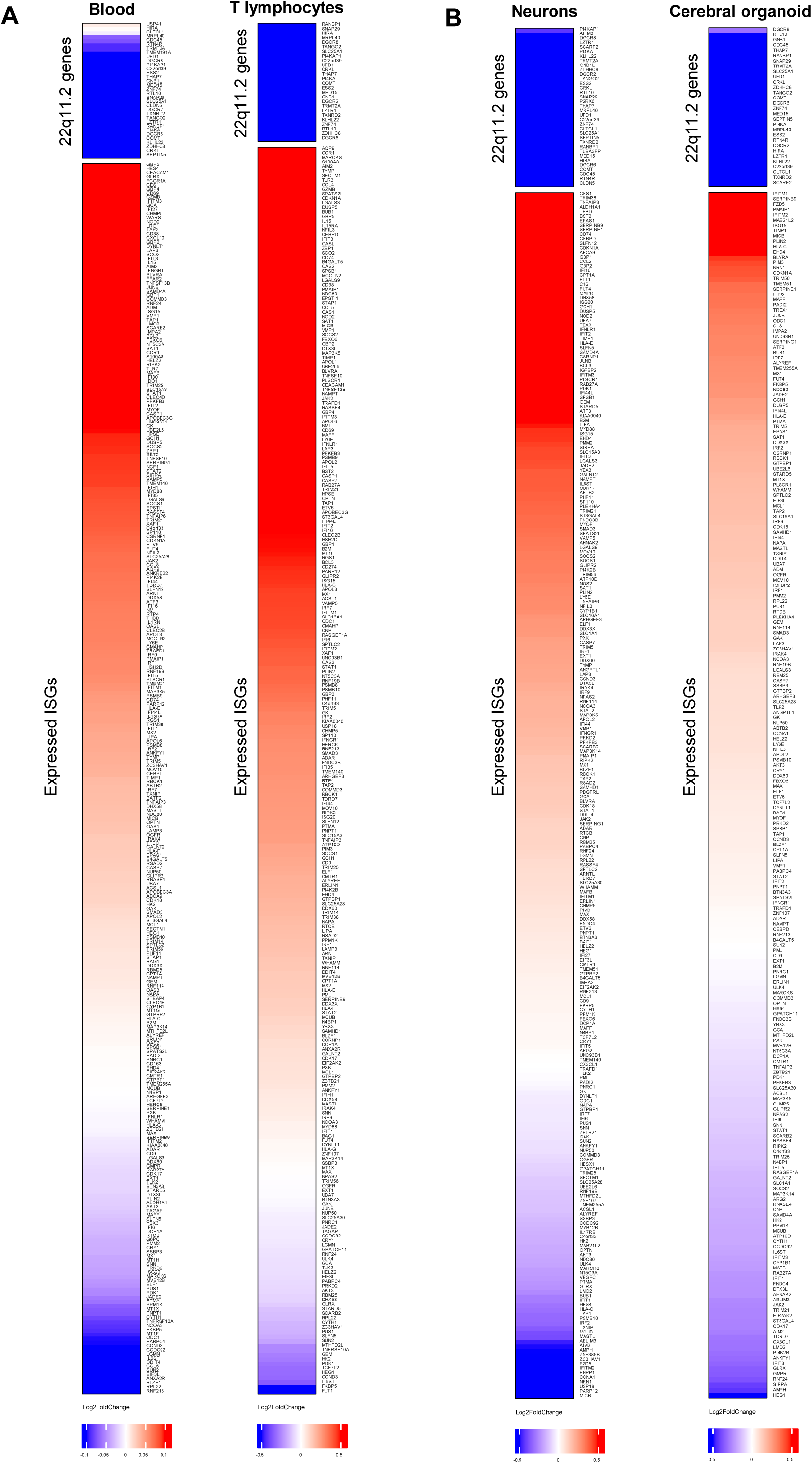
a. Heatmap showing differential gene expression of expressed 22q11.2 genes (upper panel) and expressed ISGs (lower) in blood (Lin et al. 2021) and T-lymphocytes (Raje et al. 2022). Red indicates upregulated expression in 22qDS vs control, blue indicates downregulation. b. Heatmap showing differential gene expression of expressed 22q11.2 genes (upper panel) and expressed ISGs (lower) in iPSCs-derived neurons (Lin et al. 2016) and cerebral cortex organoids differentiated from 22qDS-iPSCs (Khan et al. 2020). Red indicates upregulated expression in 22qDS vs control, blue indicates downregulation.

**SUPPLEMENTARY FIGURE 6, related to Figure 6.**
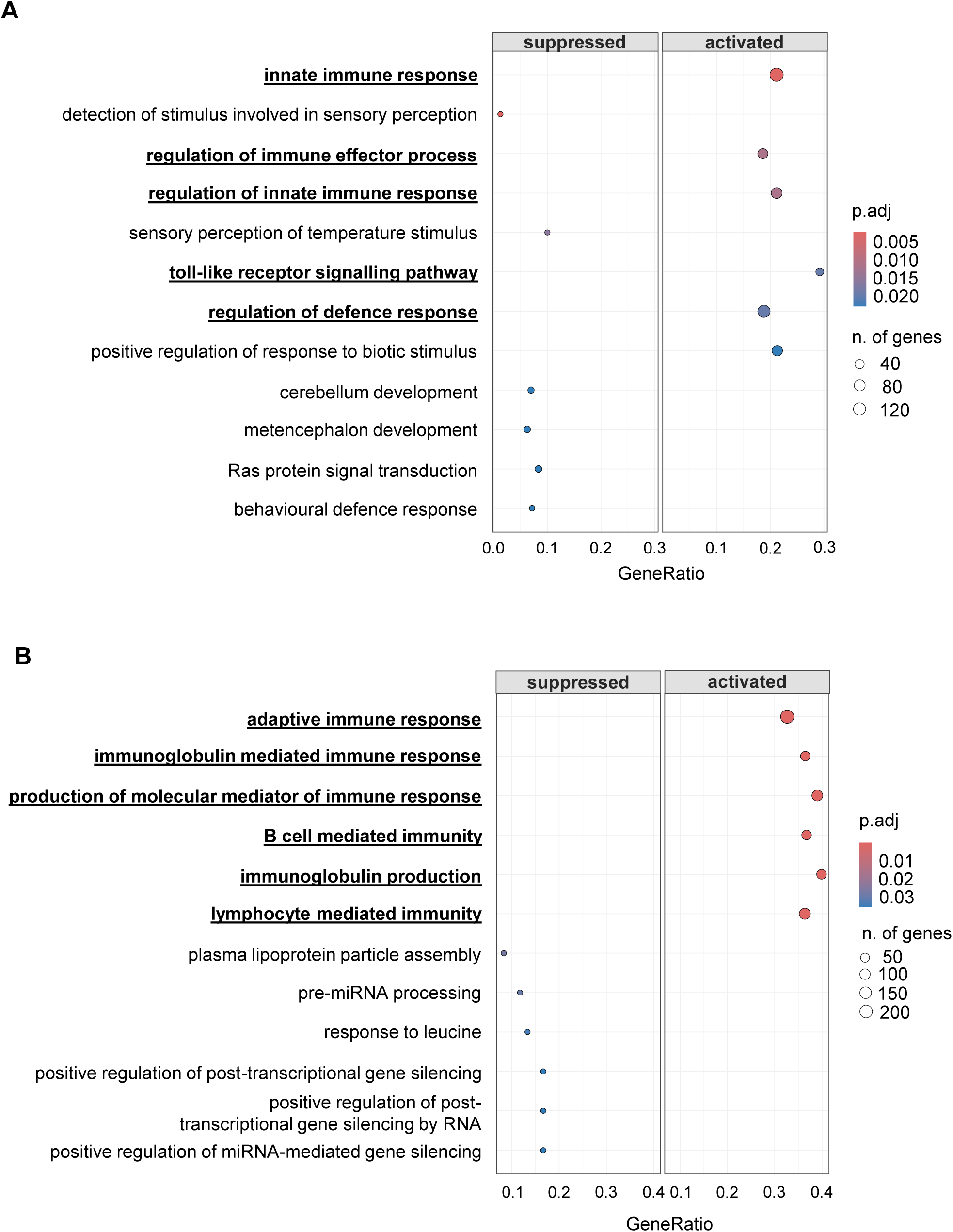
a. GSEA in 22qDS blood (n=79 patients, n=68 controls) (Lin et al. 2021) ranked by lowest adjusted p-values. Immune-related terms are in bold and underscored. b. GSEA in 22qDS T-lymphocytes (n=13 patients, n=6 controls) (Raje et al. 2022) ranked by lowest adjusted p-values. Immune-related terms are in bold and underscored.

**SUPPLEMENTARY FIGURE 7, related to Figure 6.**
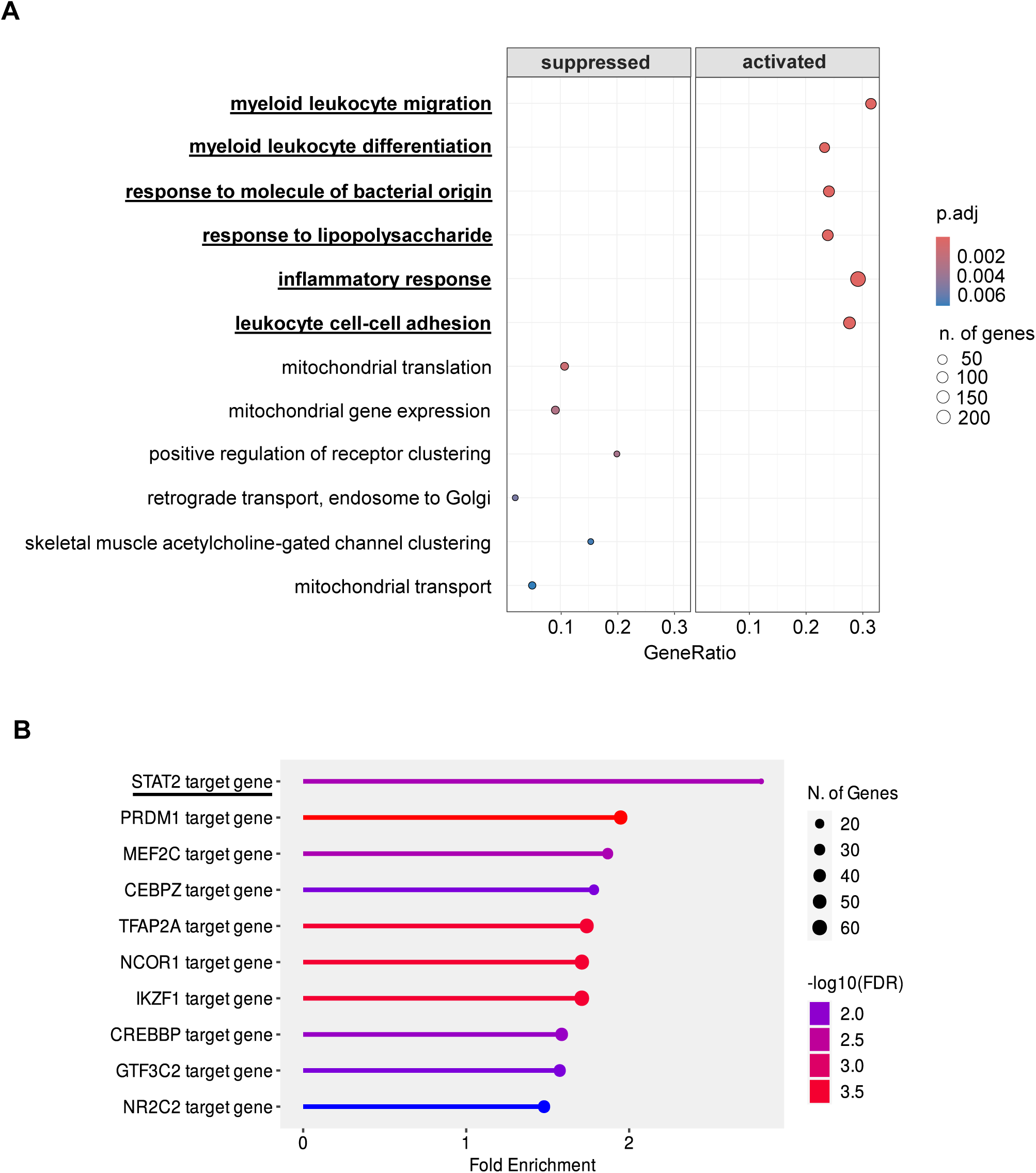
a. GSEA in 22qDS iPSCs-derived neurons (n=8 patients, n=7 controls) (Lin et al. 2016) ranked by lowest adjusted p-values. Immune-related terms are in bold and underscored. b. Transcription Factor (TF) targets network (ENCODE) obtained with upregulated genes (log2FC > 1 and p-adj < 0.05) in iPSCs-derived neurons (n=8) vs control (n=7) (Lin et al. 2016).

**SUPPLEMENTARY FIGURE 8, related to Figure 6.**
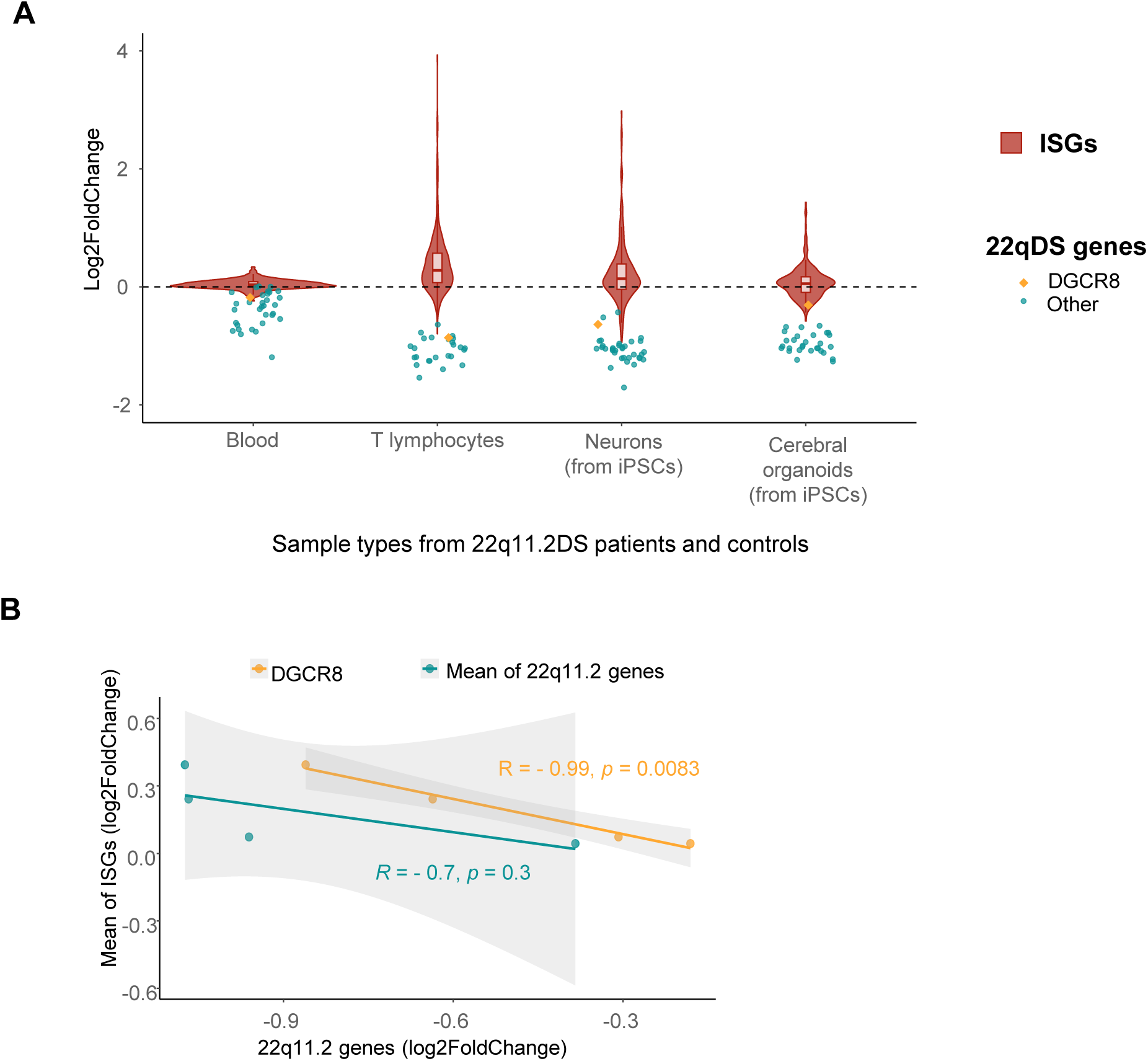
a. Differential gene expression of ISGs and 22q11.2 deleted genes in different datasets: blood (n=79 patients, n=68 controls)* (Lin et al. 2021), T-lymphocytes (n=13 patients, n=6 controls) (Raje et al. 2022), iPSCs-derived neurons (n=8 patients, n=7 controls) (Lin et al. 2016) and cerebral cortex organoids differentiated from 22qDS-iPSCs after 100 days (n=15 patients, n=14 controls) (Khan et al. 2020). The violin plot (red) represents the log2FC(22qDS/control) of each expressed ISG. Individual dots represent the log2FC (22qDS/control) of genes contained in the 22q11.2 regions (blue), including the *DGCR8* gene (yellow). *Blood dataset expression is determined by microarray and not RNA-sequencing, explaining smaller log2FCs found (Lin et al. 2021). b. Correlation between the log2FC (22qDS/control) of genes contained in the 22q11.2 region and ISGs. On the Y-axis, the log2FC of the mean of ISGs is represented. On the X-axis, the log2FC of the mean of 22q11.2 genes (blue) or DGCR8 (orange). Each dot represents one dataset, these being: blood (n=79 patients, n=68 controls) (Lin et al. 2021), T-lymphocytes (n=13 patients, n=6 controls) (Raje et al. 2022), iPSCs-derived neurons (n=8 patients, n=7 controls) (Lin et al. 2016) and cerebral cortex organoids differentiated from 22qDS-iPSCs after 100 days (n=15 patients, n=14 controls) (Khan et al. 2020).

**SUPPLEMENTARY FIGURE 9.**
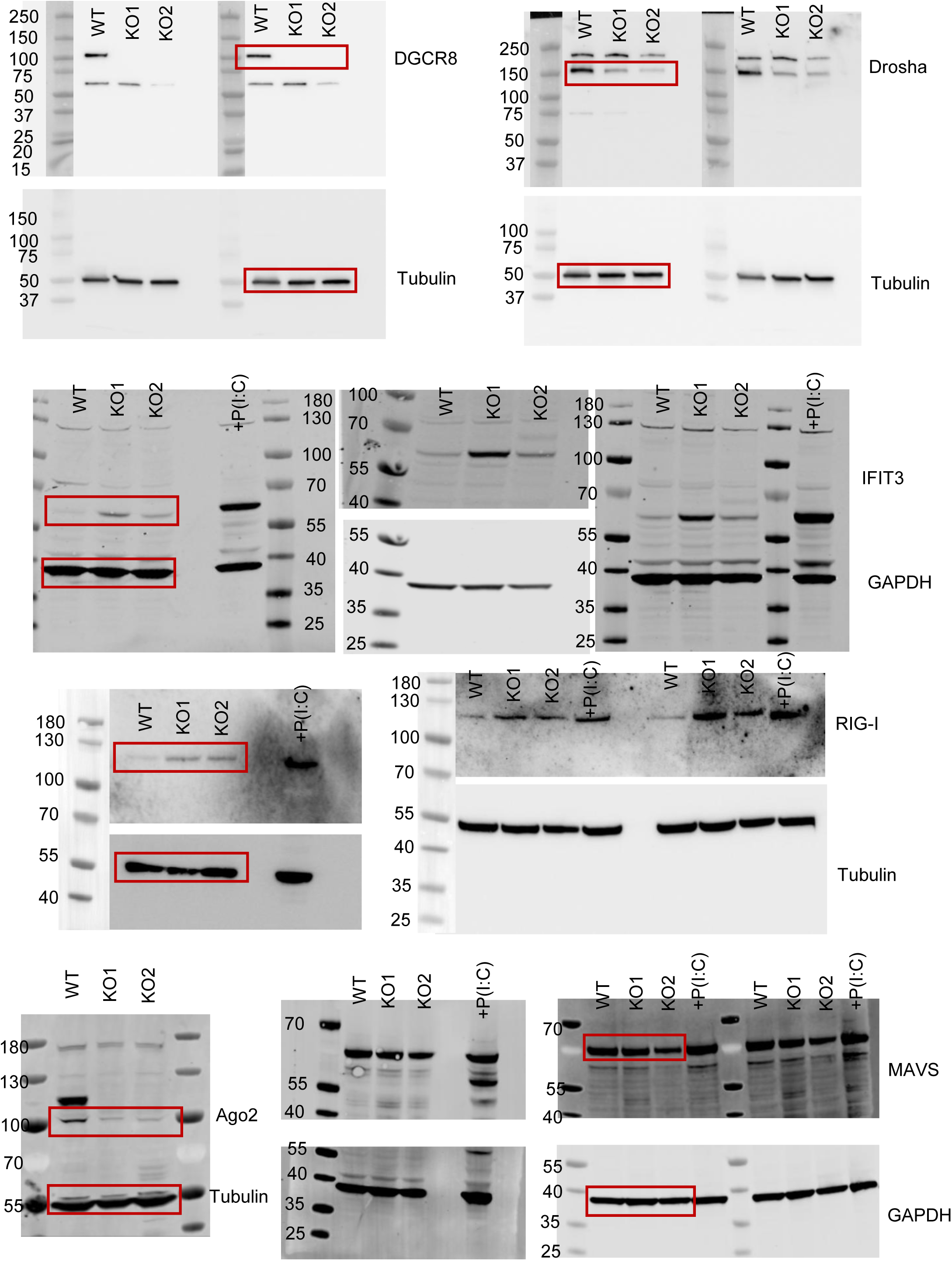
Uncropped western blots membranes corresponding to Figure 1A, Figure 1B, Figure 1H, Figure 1I, Supplementary Figure 3A and Supplementary Figure 3C.

**SUPPLEMENTARY FIGURE 10.**
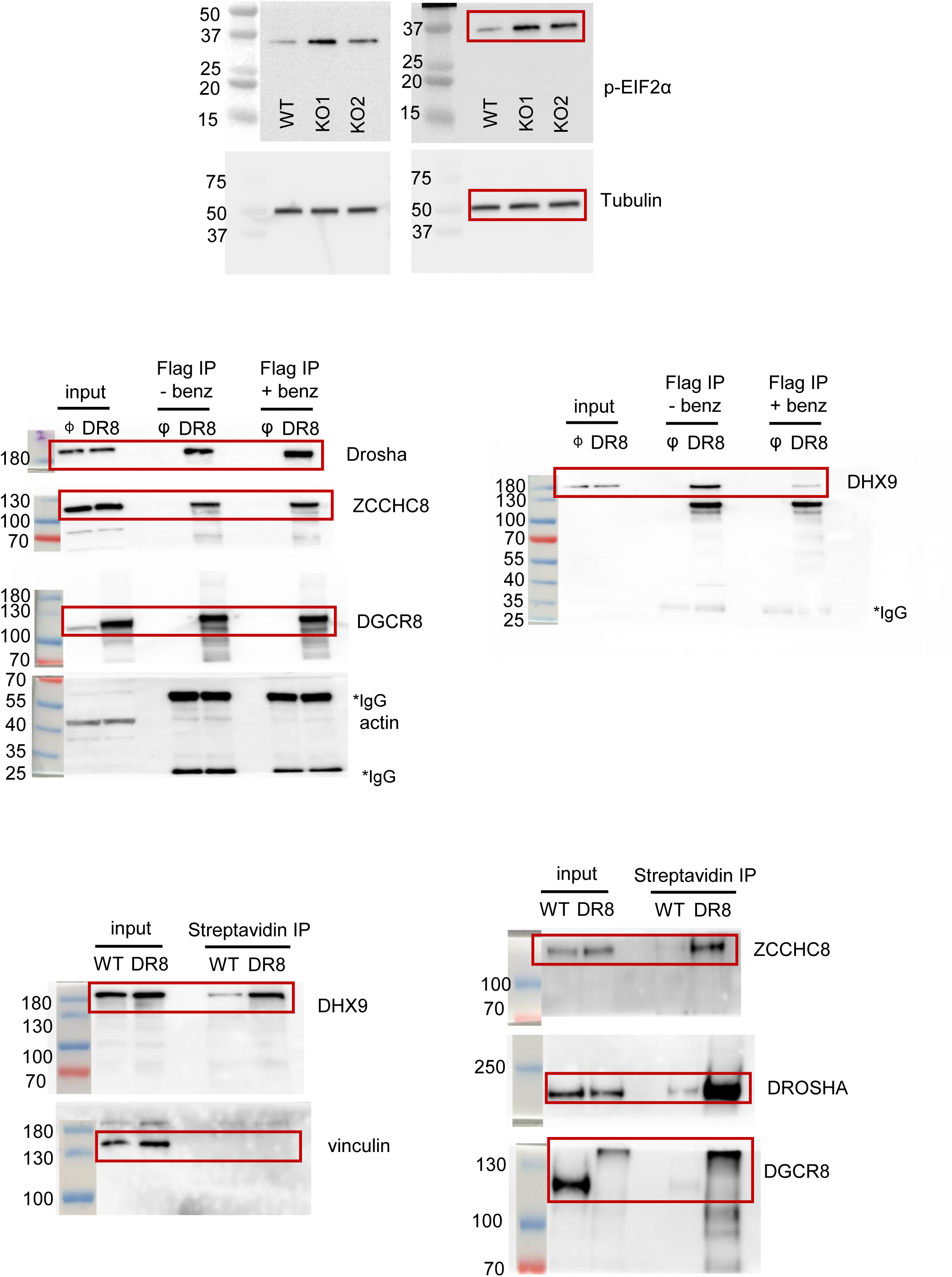
Uncropped western blots membranes corresponding to Figure 3E, Figure 5G and Figure 5H.

**SUPPLEMENTARY FIGURE 11.**
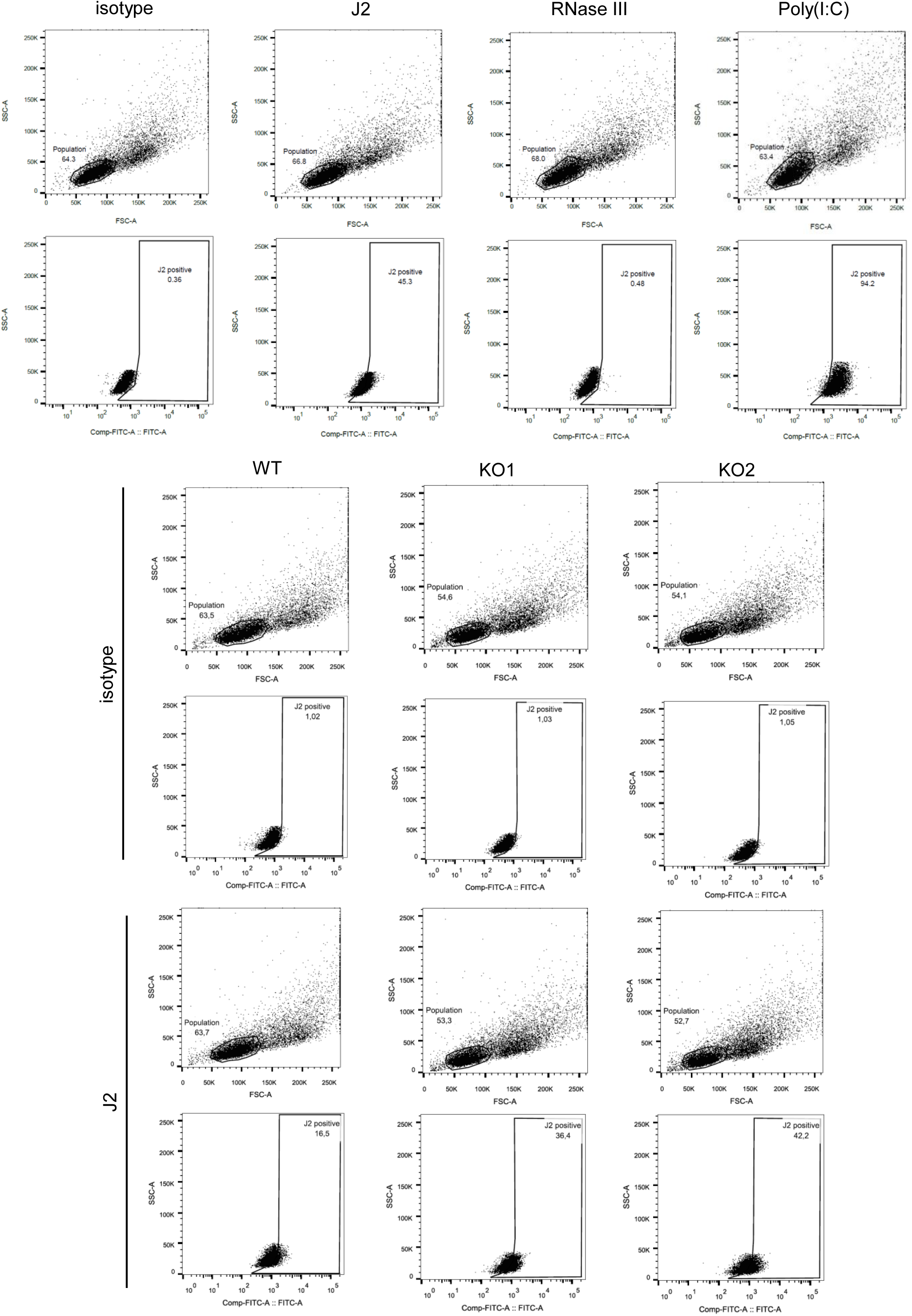
Gating strategy corresponding to Figure 3A and Figure 3C.

